# Impact of sickle cell trait hemoglobin on the intraerythrocytic transcriptional program of *Plasmodium falciparum*

**DOI:** 10.1101/2021.08.06.455439

**Authors:** Joseph W. Saelens, Jens E.V. Petersen, Elizabeth Freedman, Robert C. Moseley, Drissa Konaté, Seidina A.S. Diakité, Karim Traoré, Natalie Vance, Rick M. Fairhurst, Mahamadou Diakité, Steven B Haase, Steve M Taylor

## Abstract

Sickle-trait hemoglobin (HbAS) confers near-complete protection from severe, life-threatening falciparum malaria in African children. Despite this clear protection, the molecular mechanisms by which HbAS confers these protective phenotypes remain incompletely understood. As a forward genetic screen for aberrant parasite transcriptional responses associated with parasite neutralization in HbAS red blood cells (RBCs), we performed comparative transcriptomic analyses of *Plasmodium falciparum* in normal (HbAA) and HbAS erythrocytes during both *in vitro* cultivation of reference parasite strains and naturally-occurring *P. falciparum* infections in Malian children with HbAA or HbAS. During *in vitro* cultivation, parasites matured normally in HbAS RBCs, and the temporal expression was largely unperturbed of the highly ordered transcriptional program that underlies the parasite’s maturation throughout the intraerythrocytic development cycle (IDC). However, differential expression analysis identified hundreds of transcripts aberrantly expressed in HbAS, largely occurring late in the IDC. Surprisingly, transcripts encoding members of the Maurer’s clefts were overexpressed in HbAS despite impaired parasite protein export in these RBCs, while parasites in HbAS RBCs underexpressed transcripts associated with the endoplasmic reticulum and those encoding serine repeat antigen proteases that promote parasite egress. Analyses of *P. falciparum* transcriptomes from 32 children with uncomplicated malaria identified stage-specific differential expression: among infections composed of ring-stage parasites, only cyclophilin 19B was underexpressed in children with HbAS, while trophozoite-stage infections identified a range of differentially-expressed transcripts, including downregulation in HbAS of several transcripts associated with severe malaria in collateral studies. Collectively, our comparative transcriptomic screen *in vitro* and *in vivo* indicates that *P. falciparum* adapts to HbAS by altering its protein chaperone and folding machinery, oxidative stress response, and protein export machinery. Because HbAS consistently protects from severe *P. falciparum*, modulation of these responses may offer avenues by which to neutralize *P. falciparum* parasites.

**Importance:** Sickle-trait hemoglobin (HbAS) confers near-complete protection from severe, life-threatening malaria, yet the molecular mechanisms that underlie HbAS protection from severe malaria remain incompletely understood. Here, we use RNA-seq to measure the impact of HbAS on the blood stage transcriptome of *Plasmodium falciparum* in *in vitro* time series experiments and *in vivo* samples from natural infections. Our i*n vitro* time series data reveal that, during its blood stage, *P. falciparum’s* gene expression in HbAS is impacted primarily through alterations in the abundance of gene products as opposed to variations in the timing of gene expression. Collectively, our *in vitro* and *in vivo* data indicate that *P. falciparum* adapts to HbAS by altering its protein chaperone and folding machinery, oxidative stress response, and protein export machinery. Due to the persistent association of HbAS and protection from severe disease, these processes that are modified in HbAS may offer strategies to neutralize *P. falciparum*.

## Introduction

In malaria-endemic regions across Africa, red blood cell (RBC) variants are highly prevalent and can confer significant protection against severe malaria (1). Heterozygosity for sickle hemoglobin (Hb) provides one striking example of the protection afforded by an RBC variant: sickle-cell trait (HbAS) reduces the risk of severe, life-threatening malaria by over 90% (2). The functional underpinnings of this extensive protection afforded by HbAS against severe disease hold potential for uncovering mechanisms of parasite pathogenesis that can be exploited by future interventions.

Candidate protective mechanisms reported thus far are diverse and provide evidence that HbAS disrupts mediators of pathogenesis and the parasite life cycle in infected RBCs (iRBCs). Parasite maturation may be inhibited by either low oxygen tension (3–6) or host miRNAs that are enriched in HbSS and HbAS RBCs (7), though others have reported equivalent growth in HbAA and HbAS iRBCs in the same conditions (8, 9). Multiple lines of evidence support a mechanism in which cytoadherence, a central property of *P. falciparum’s* pathogenesis, is reduced in HbAS iRBCs. HbAS reduces both the display on the iRBC surface of *P. falciparum* erythrocyte membrane protein-1 (PfEMP-1) as well as knob density (10). This is supported by the observation that HbAS impairs export of proteins including PfEMP-1 to the iRBC surface via Maurer’s clefts, the Golgi-like organelles in the iRBC cytosol that sort and traffic proteins (11). The attenuated export of surface proteins and concomitant reduction in cytoadherence would compromise the capacity of HbAS iRBCs to sequester in the microvasculature endothelium, thereby diminishing endothelial activation - as has been observed *in vitro* (12) – as well as subsequent events in the microvasculature that can lead to severe disease *in vivo* (13). These downstream phenotypes of HbAS iRBCs are the result of incompletely understood upstream parasite responses to the altered RBC environment. The natural protection against severe falciparum malaria conferred by HbAS provides an opportunity to identify parasite mechanisms that are neutralized and therefore critical to pathogenesis. In this study, we use RNA-seq to perform an unbiased comparative transcriptomic survey of the *P. falciparum* transcriptional changes that occur in HbAS iRBCs. Owing to the reduced cytoadherence observed in HbAS iRBCs, we hypothesized that gene products functioning in cytoadherence would be downregulated in HbAS. Therefore, we investigated parasite transcription over the IDC *in vitro* in tightly synchronized parasite populations in HbAA and HbAS iRBCs as well as in freshly-collected parasites from Malian children with uncomplicated malaria with HbAA or HbAS.

## Results

### Parasite maturation and transcriptome characteristics

We interrogated two reference *Plasmodium falciparum* strains that originated where the sickle allele of ß-globin is common (14): 3D7 from West Africa (15) and FUP from Uganda (16). Based on morphology assessed by microscopy, parasites matured similarly in HbAA and HbAS iRBCs for both 3D7 and FUP (**Figure 1A** and data at https://github.com/duke-malaria-collaboratory/Sickle-trait-RNAseq/tree/master/_data/in_vitro/DESeq2). 3D7 parasites in HbAA and HbAS iRBCs transitioned from rings to trophozoites at 27hpi in 3D7 and by 21hpi in FUP.

**Fig 1.**
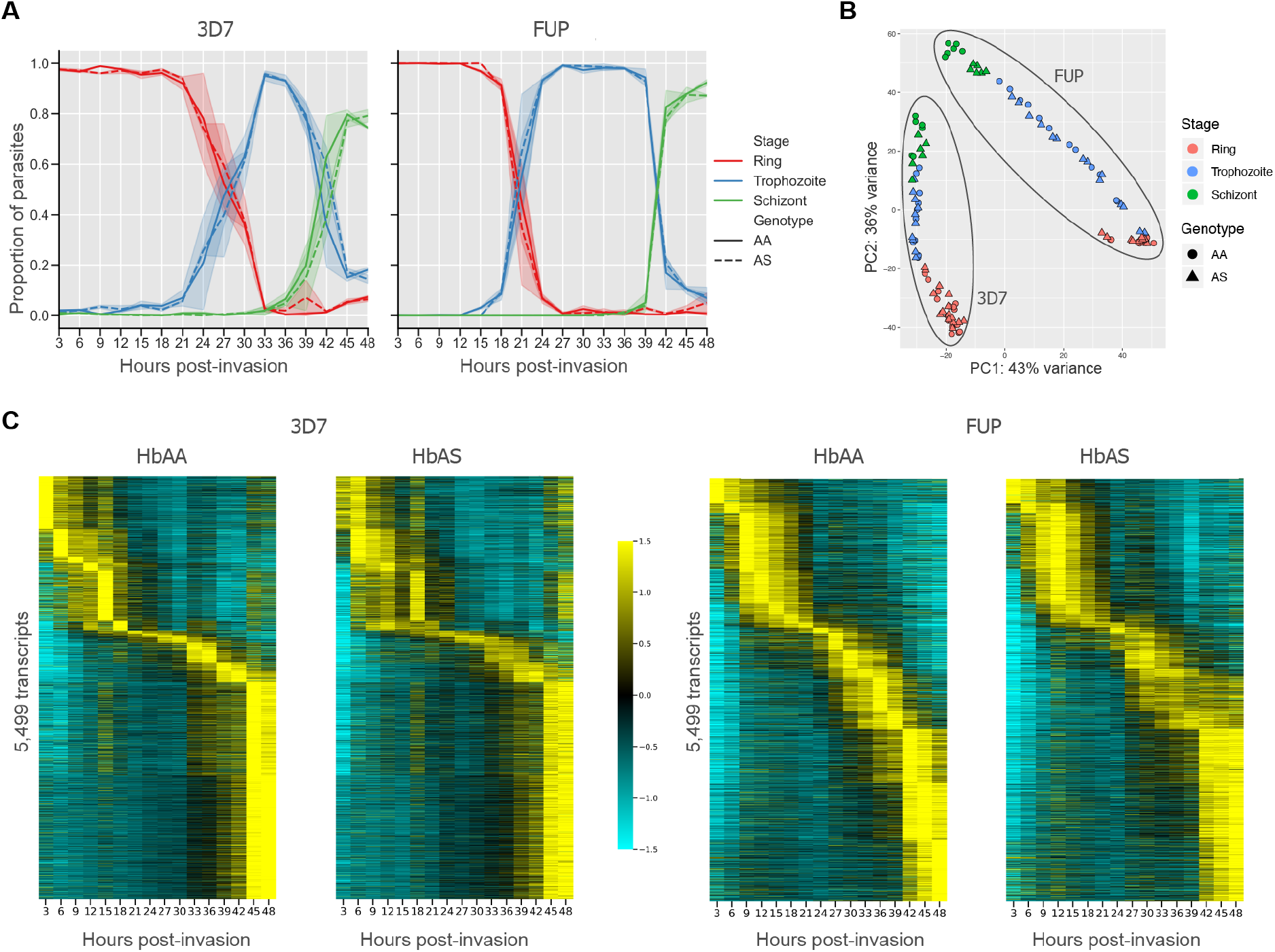
*P. falciparum* maturation and overall transcriptional program are conserved in HbAS iRBCs. **(A)** The relative abundance of ring, trophozoite, and schizont stage parasites from light microscopy readings in HbAA (solid line) and HbAS (dotted line) iRBCs. 3D7 and FUP parasites mature at similar rates in HbAA and HbAS iRBCs. Counts are based on 500 parasites from each sample per time point and were assessed by a reader masked to timepoint and RBC type. **(B)** Principal component analysis of transcript expression following variance-stabilizing transformation based on the 500 transcripts with the highest variance across all samples. Samples cluster according to parasite strain and developmental stage. **(C)** Expression of 5,499 transcripts passing thresholds in the ASM276v2 transcriptome in 3D7 and FUP isolates. In both 3D7 and FUP, the ordering of expression is largely conserved between HbAA and HbAS samples. Transcripts are ordered vertically by the peak expression time in HbAA RBCs for each parasite strain. The columns along the *x*-axis depict the normalized expression values of each transcript per time point. Expression values were averaged across replicates, and are normalized around the mean for each transcript over the time series and depicted as a *z* score of standard deviations from the mean. Values range between −1.5 (cyan) and 1.5 (yellow). Scripts for RNA-seq read preparation and quantification are available on our GitHub page.

Principal component analysis (PCA) based on transcript counts clustered samples by parasite line and developmental stage (**Figure 1B**). Notably, the principal components that explain the vast majority of the transcriptional variance between samples did not separate parasite transcriptomes in HbAS iRBCs from those in HbAA iRBCs. Similarly, visualization of normalized expression of 5,499 parasite transcripts suggests that the progression of *P. falciparum’s* IDC transcriptional program remains largely intact in HbAS iRBCs (**Figure 1C**). These data indicate both that the parasite populations in each sample were highly synchronous and that HbAS does not broadly disrupt *P. falciparum’s* transcriptional program during the IDC.

### Transcriptional synchrony of parasites in HbAS RBCs

*P. falciparum* maturation in HbAS iRBCs has previously been described to be delayed compared to parasites in HbAA iRBCs (6). To assess parasite maturation at a finer scale, we computed, for the subset of transcripts that had a single transcription peak, the temporal shift of each transcript’s expression peak in HbAS iRBCs compared to HbAA (**Figure 2A, Figure S2**). The distribution of peak shifts between HbAA and HbAS over the entire time course was centered around zero for both 3D7 and FUP (**Figure 2B, Figure S3**), with slight but nonsignificant mean peak shift delays in HbAS RBCs for both 3D7 (−1.62 hours) and FUP (−0.91 hours) parasites (**Figures S3**).

**Fig. 2.**
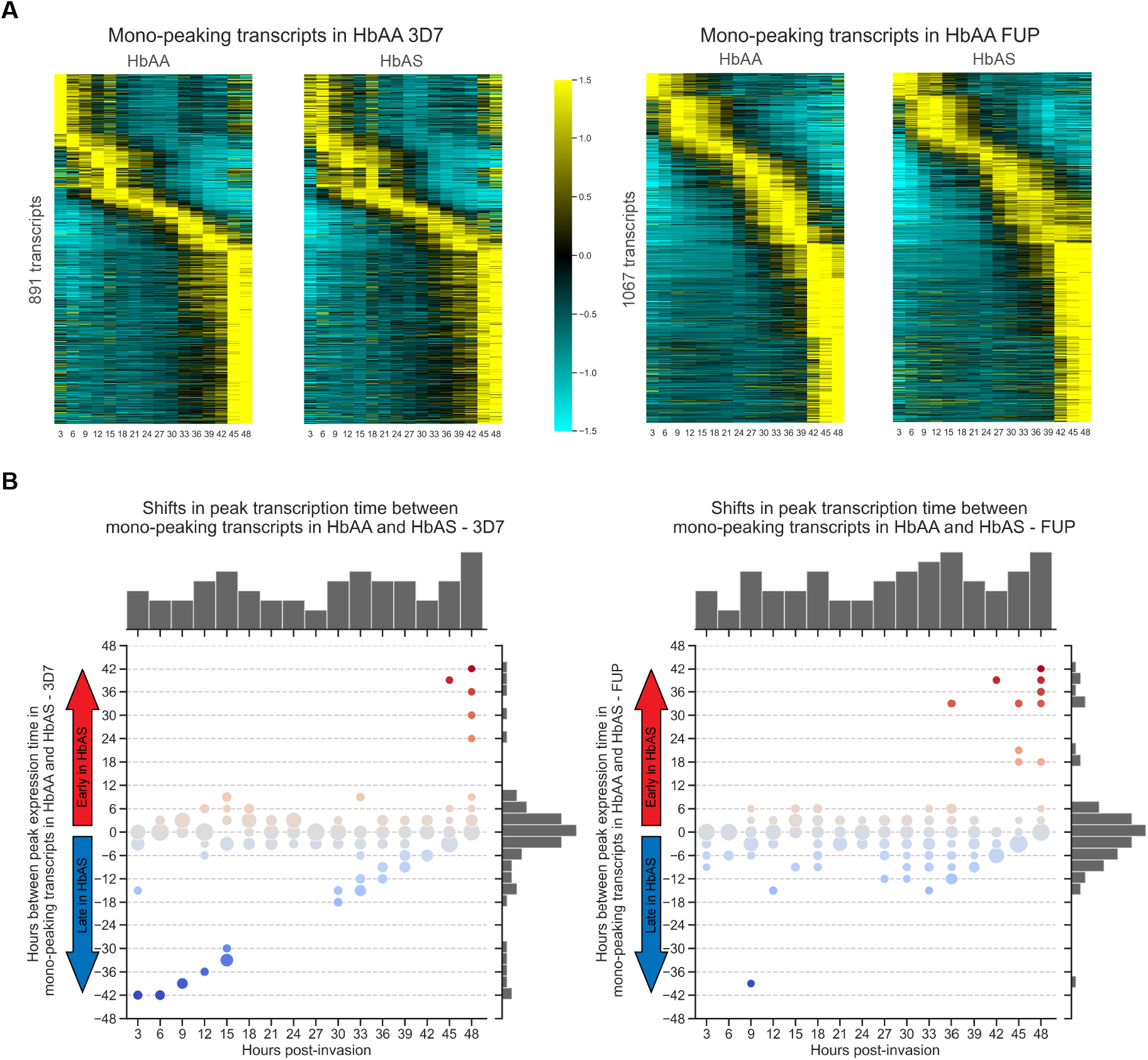
Shifts in peak expression time between HbAA and HbAS. **(A)** Peak expression time of the subsets of mono-peaking parasite transcripts in HbAA and HbAS iRBCs for 3D7 (left pair) or FUP (right pair) parasites. For each pair, transcripts are ordered vertically by the peak expression time in HbAA iRBCs. In both 3D7 and FUP, the ordering of expression is largely conserved between HbAA and HbAS samples. The columns along the *x*-axis depict the normalized expression values of each transcript per time point. Expression values were averaged across replicates, and are normalized around the mean for each transcript over the time series and depicted as a *z* score of standard deviations from the mean. Values range between −1.5 (cyan) and 1.5 (yellow). **(B)** Temporal shifts for peak expression timepoint for the subset of mono-peaking parasite transcripts in HbAA compared with HbAS iRBCs for 3D7 (left) or FUP (right) parasites. For each transcript, the time between its expression peak in parasites in HbAA and HbAS iRBCs was calculated and averaged across ß-globin replicates for each strain. Each *x*-axis category indicates the number of transcripts that normally peak at this timepoint in HbAA iRBCs, and the y-values indicate the distributions of the shifts of the peaks of these transcripts in HbAS iRBCs. Circles are sized relative to the number of transcripts with that y-value. Top histogram indicates the number of parasite transcripts that peak in HbAA iRBCs at each *x*-value timepoint, and right histogram indicates the number of transcripts with peak shifts of each *y*-value. The distribution of peak shifts (displayed along the right *y*-axis) is centered around zero, indicating that most transcripts peak at the same time in parasites growing in HbAA and HbAS iRBCs. See Jupyter Notebook “expressionPeakChanges.ipynb” on our GitHub for full workflow.

### Differentially expressed transcripts identified in HbAS *in vitro*

HbAS impacts *P. falciparum’s* transcriptional program in the IDC at varying time points in 3D7 and FUP isolates (**Figures 3A** and **3B** and data at https://github.com/duke-malaria-collaboratory/Sickle-trait-RNAseq/tree/master/_data/in_vitro/DESeq2), though in both strains we observed an absence of differential expression through much of the ring stage, with transcriptional divergence apparent in the mid- to late-trophozoite stage that accelerated as parasites entered the schizont stage (**Figure 3B**).

**Fig 3.**
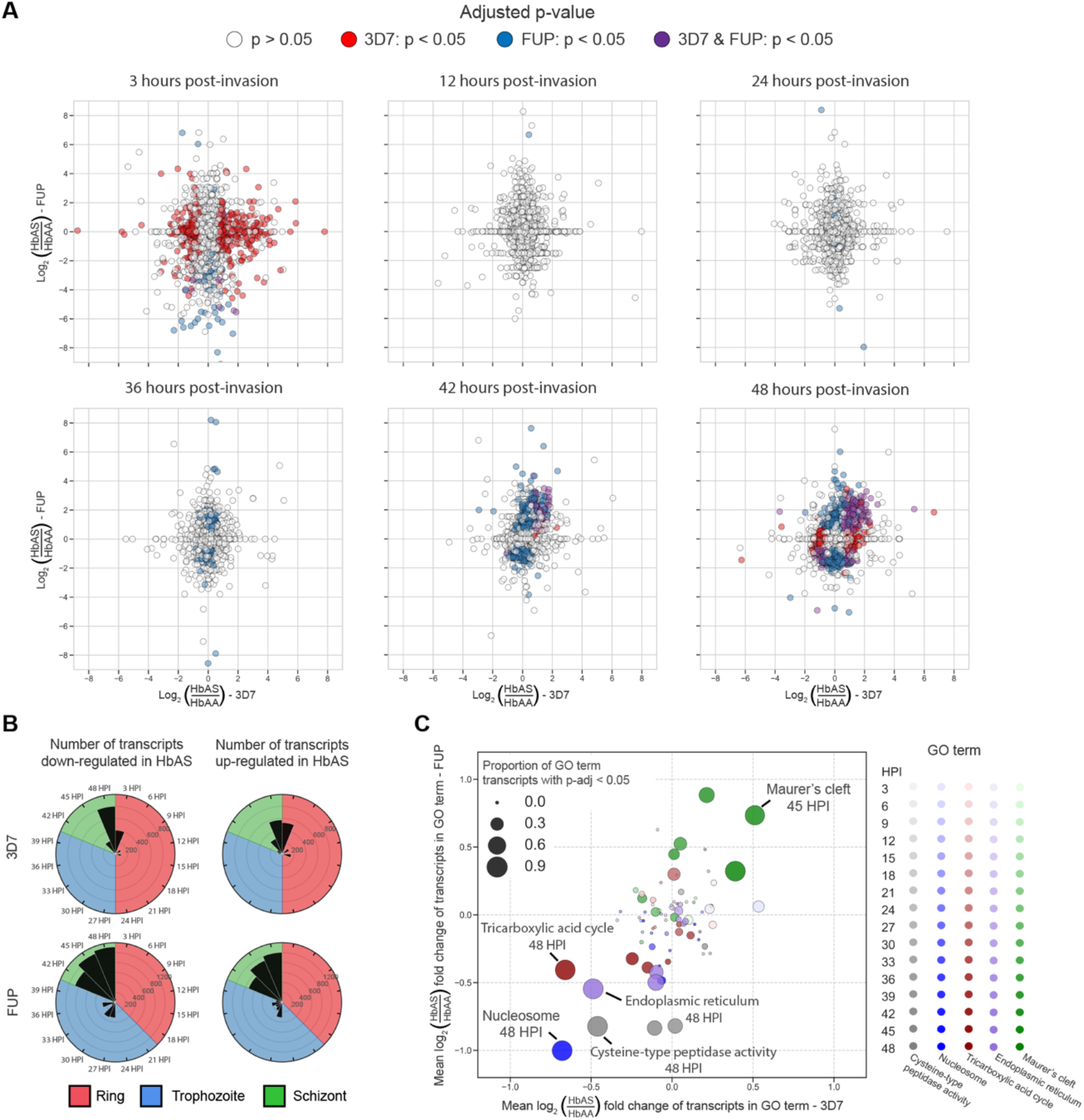
Differential expression analysis reveals parasite transcript expression in HbAS iRBCs diverges from HbAA parasites primarily in the schizont stage. **(A)** Scatterplots at 3-, 12-, 24-, 36-, 42-, and 48 hours post-invasion of the log2 fold-change of individual transcripts for HbAS parasites compared to HbAA parasites (hpi) in 3D7 (*x*-axes) and FUP (*y*-axes) parasites. Points in the upper right and lower left quadrants signify transcripts that are differentially expressed in the same direction between parasite strains, i.e. those in the upper right quadrant are upregulated in both 3D7 and FUP in HbAS iRBCs, and those in the lower left quadrant are downregulated in both 3D7 and FUP in HbAS iRBCs. Point color indicates the statistical significance of DESeq2 differential expression analysis for each transcript: white points have an adjusted p-value > 0.05 in both 3D7 and FUP, red points have an adjusted p-value < 0.05 in 3D7 but not FUP, blue points have an adjusted p-value < 0.05 in FUP but not 3D7, and purple points have an adjusted p-value < 0.05 in both 3D7 and FUP. **(B)** Counts of transcripts in 3D7 (top) and FUP (bottom) classified as downregulated (left column) or upregulated (right column) at a significance level of adjusted p-value < 0.05 at each timepoint, from 3hpi to 48hpi. Shading of each plot is colored according to the predominant stage of the parasites at each timepoint: ring (red), trophozoite (blue), or schizont (green). There were minimal shared differentially expressed transcripts until 39 hpi, after which 966 transcripts were differentially expressed in both 3D7 and FUP (493 downregulated and 460 upregulated). **(C)** Scatterplot of transcript mean fold changes within gene ontology (GO) terms that are enriched for differentially expressed transcripts in HbAS iRBCs in both 3D7 (*x*-axis) and FUP (*y*-axis) parasites. Color indicates GO term, shading indicates hpi (darker = later hpi), and size of point indicates the proportion of transcripts within the GO term that are differentially expressed at an adjusted p-value <0.05. Significant expression changes of GO terms occurred at later time-points during the schizonts stage, including upregulation of Maurer’s cleft transcripts in HbAS iRBCs and concurrent downregulation of transcripts involved in the endoplasmic reticulum, the tricarboxylic acid cycle, cysteine-type peptidase activity, and nucleosomes.

We identified the transcripts that were differentially expressed in the same direction in both 3D7 and FUP at each time point (**Figure 3A**, purple dots, **Supplementary Table 1**), which indicated a consistent late-stage effect of HbAS on transcriptional phenotypes across parasite strains (**Figure 3B**). In a gene ontology (GO) enrichment analysis of this shared set of transcripts (17) (**Figure 3C**), the only GO term that was enriched for upregulated transcripts in both 3D7 and FUP in HbAS iRBCs at any time point in our time series experiments was that for Maurer’s clefts (**Figure 4A**), which are parasite-derived membranous structures that sort and traffic proteins and in HbAS iRBCs are both dysmorphic and dysfunctional (8). Several critical components of Maurer’s clefts, including the membrane associated histidine-rich protein 1 (MAHRP1, PF3D7_1370300), and skeleton-binding protein 1 (SBP1, PF3D7_0501300), are encoded by transcripts that were among the most significantly upregulated in HbAS RBCs.

**Fig 4.**
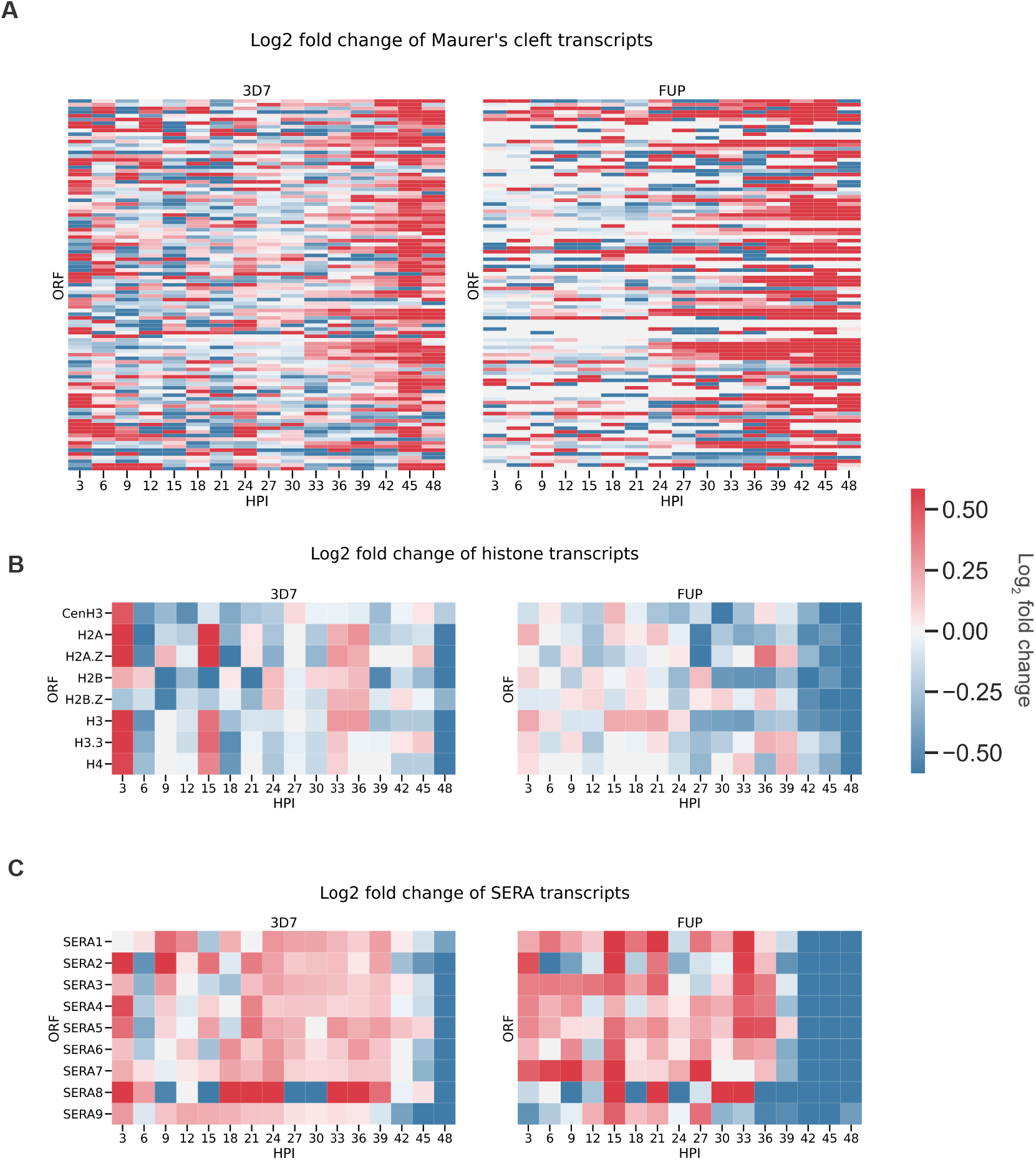
Differential expression of Maurer’s cleft, histone, and SERA transcripts in HbAS RBCs. (A) Heatmaps of log_2_ fold changes between HbAS and HbAA of Maurer’s cleft transcripts in 3D7 and FUP parasites. Transcripts IDs were extracted from the Maurer’s cleft GO term (GO:0020036) (B) Heatmaps of log_2_ fold changes between HbAS and HbAA of histone transcripts in 3D7 and FUP parasites. Transcripts IDs were extracted from the nuclecome GO term (GO:0000786). (C) Heatmaps of log_2_ fold changes between HbAS and HbAA of SERA transcripts in 3D7 and FUP parasites. Color bar is restricted to log2 fold changes considered significant in this study (|log_2_(HbAS/HbAA)| > log_2_(1.5)).

The most downregulated GO term in both 3D7 and FUP in HbAS iRBCs was that of the nucleosome (GO:0000786), which comprises eight transcripts that encode the core histones and histone variants in *P. falciparum*. In both 3D7 and FUP growing in HbAS RBCs, all histone transcripts were downregulated at 48 hpi (**Figure 4B**). Similarly, downregulated transcripts were enriched in the TCA cycle (GO:0006099) and the endoplasmic reticulum (ER) (GO:0005783) at late time points in 3D7 and FUP in HbAS iRBCs. The ER GO term encompasses a wide range of proteins, including heat shock proteins, phosphatases, peptidyl-prolyl cis-trans isomerases, and plasmepsins. Of 53 transcripts in the ER GO term, 30 transcripts were downregulated in both strains in HbAS iRBCs between 45-48 hpi, suggesting a broad inhibition of protein export in late-stage parasites.

Among Molecular Function GO terms, downregulated transcripts were enriched only in that of cysteine-type peptidase activity (GO:0008234), which primarily comprises the serine repeat antigen (SERA) proteases that play a pivotal role enabling parasite egress from the iRBC. SERA2-7 were significantly downregulated in 3D7 and FUP parasites in HbAS iRBCs at late time points (**Figure 4C**), suggesting an impairment of merozoite egress that may help to explain the lower parasite densities observed in sickle-trait children with malaria (18).

### Temporal dysregulation of transcript expression in HbAS iRBCs

To identify the transcripts that are temporally dysregulated, we analyzed our time series data using dynamic time warping (DTW) (**Figure 5A**), which determines between two series the optimal alignment of a transcript’s expression and assigns a score as a measure of similarity (low score, **Figure 5B – top row**) or difference (high score, **Figure 5B – bottom row**). Overall, DTW scores comparing transcript temporal expression between HbAA and HbAS RBCs were close to normally-distributed with a positive skew in 3D7 and in FUP (**Figure 5A**). Between strains, mean transcript DTW scores comparing HbAA versus HbAS time series were positively linearly correlated when computed for 3D7 or FUP (r = 0.524, p-value < 0.0001), indicating a conserved impact on transcript timing across strains (**Figure S9A**).

**Fig 5.**
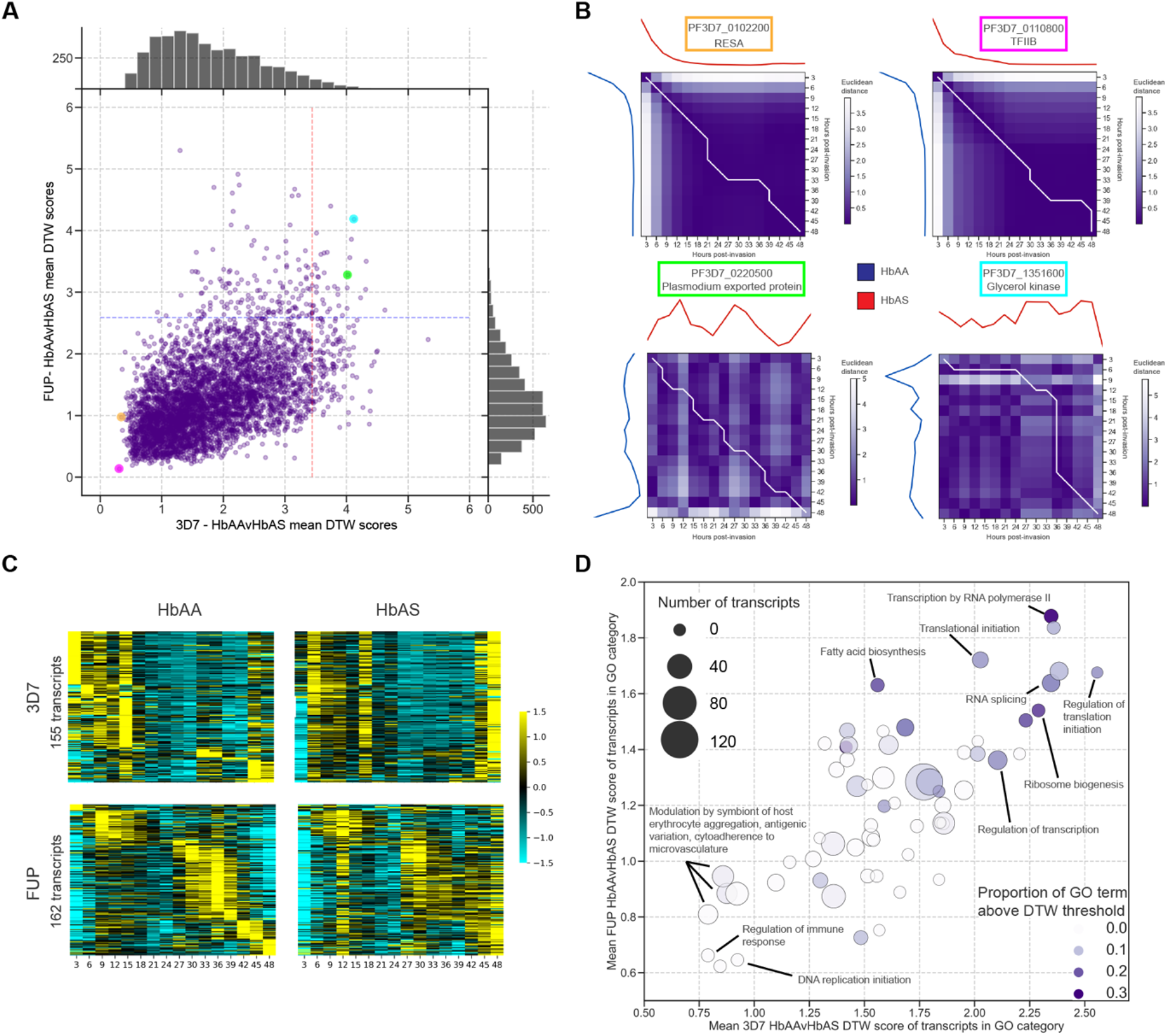
HbAS impacts the temporal expression of transcripts governing gene regulation. **(A)** Scatterplot of dynamic time warping (DTW) scores in time series data of individual transcripts in HbAA compared to HbAS samples in 3D7 (*x*-axis) and FUP (*y*-axis) parasites. For each strain, scores were computed pairwise between HbAA and HbAS samples and averaged. Transcripts with dynamic time warping scores greater than two standard deviations from the overall mean (points to the right of the red line for 3D7, and points above the blue line for FUP) were considered temporally dysregulated. Histograms indicate overall distribution of transcript DTW scores in either 3D7 (top) or FUP (right). **(B)** The Euclidean distance scores and warping path of selected transcripts that correspond to the highlighted circles in **Figure 5A**. Mean-normalized transcript expression is plotted along the *x*- and *y*-axes: that for HbAA iRBCs is plotted from top to bottom along the left *y*-axis, and for HbAS iRBCs is plotted from left to right along the upper *x*-axis. The heatmap matrix is colored according to the Euclidean distance between the mean-normalized expression values between HbAA and HbAS at the corresponding time points. Dynamic time warping scores are cumulatively computed along the warping path (white line). Transcripts for the ring-infected erythrocyte surface antigen (RESA, PF3D7_010220) and the putative transcription factor TFIIB (PF3D7_0110800) (top) had very low DTW scores between HbAA and HbAS samples in both strains, while those for the Plasmodium exported protein (hyp2, PF3D7_0220500) and a glycerol kinase (PF3D7_1351600) had high DTW scores indicating dissimilar transcriptional profiles between HbAA and HbAS RBCs (bottom). **(C)** Heatmaps of mean-normalized expression of the 155 (3D7, top) and 162 (FUP, bottom) transcripts with DTW scores exceeding two standard deviations from the mean score in HbAA (left) and HbAS (right) iRBCs (points beyond the dashed lines in **Figure 5A**). Transcripts for each parasite strain are order vertically by their peak expression time in HbAA iRBCs. The ordering of peak expression in HbAS samples is not conserved in temporally dysregulated transcripts identified by DTW. Transcripts are ordered by peak expression time in HbAA samples and displayed as a *z* score of standard deviations from the mean. **(D)** Scatterplot of mean DTW scores within GO terms that are enriched for temporally-dysregulated transcripts in HbAS iRBCs in both 3D7 (*x*-axis) and FUP (*y*-axis) parasites. Color indicates GO term, shading indicates hpi (darker = later hpi), and size of point indicates the proportion of transcripts within the GO term that are temporally-dysregulated as defined by a DTW score greater than 2 standard deviations from the overall mean of DTW scores (points beyond the dashed lines in **Figure 5A**). Transcripts involved in gene regulation processes demonstrate higher DTW scores. See Jupyter Notebook “DTW.ipynb” on our GitHub for full workflow.

We observed in HbAS iRBCs 155 temporally-dysregulated transcripts in 3D7 and 162 in FUP (**Figure 5C**). In contrast to the generally unperturbed expression patterns of the overall transcriptome (**Figure 1C**), these subsets of transcripts demonstrate aberrant expression profiles in HbAS iRBCs (**Figure 5C**). To identify which biological processes were most dysregulated in their temporal expression, transcripts were grouped by GO terms and, for each term, the mean DTW score was calculated between parasites in HbAA and HbAS. We observed that transcripts associated with antigenic variation and cytoadhesion (i.e., the variant surface antigens) were among the most conserved in the timing of their expression (**Figure 5D**). In contrast, the highest DTW scores were recorded for transcripts involved in transcription by RNA polymerase II (GO:0006366), RNA splicing (GO:0008380), regulation of translational initiation (GO:0006446), and ribosome biogenesis (GO:0042254), which collectively suggest that HbAS may have post-transcriptional consequences beyond what can be observed in our data.

We focused more closely on ring-stage expression by screening with the Temporal Alignment Kendall-Tau (TAKT) algorithm, which measures the overall similarity and temporal shift between two curves (19), and then computing for these candidate transcripts DTW scores. Using this approach in ring-stage parasites, we identified 9 transcripts in 3D7 and 11 in FUP that were dysregulated between parasites in HbAA and HbAS (**Figure S10**), though none of these were shared between the two strains.

### *In vivo* transcriptomes of *P. falciparum* in children with HbAS

We next compared parasite transcriptomes between 16 uncomplicated *P. falciparum* episodes in HbAS children matched to 16 episodes in HbAA children, all collected prospectively in Kenieroba, Mali. We first used our time-series data to identify stage-specific ring (n=7 transcripts), trophozoite (n=4), and schizont (n=6) transcripts (**Figure S12**), and, using the expression levels of these 17 transcripts as stage-markers, we classified 24 infections as ring-stage and 12 as trophozoite stage (**Figure 6A, Figure S13**). PCA and hierarchical clustering supported these classifications (**Figure 6B**). Late-stage trophozoites are rare in circulation (20), so we re-estimated the stage of each infection using an independent published mixture model that assigns parasite stage at a finer resolution, and our trophozoite classifications were categorized as early trophozoites (21). Overall, the stage classifications from this second method confirmed initial findings (**Figure S13C**).

**Fig 6.**
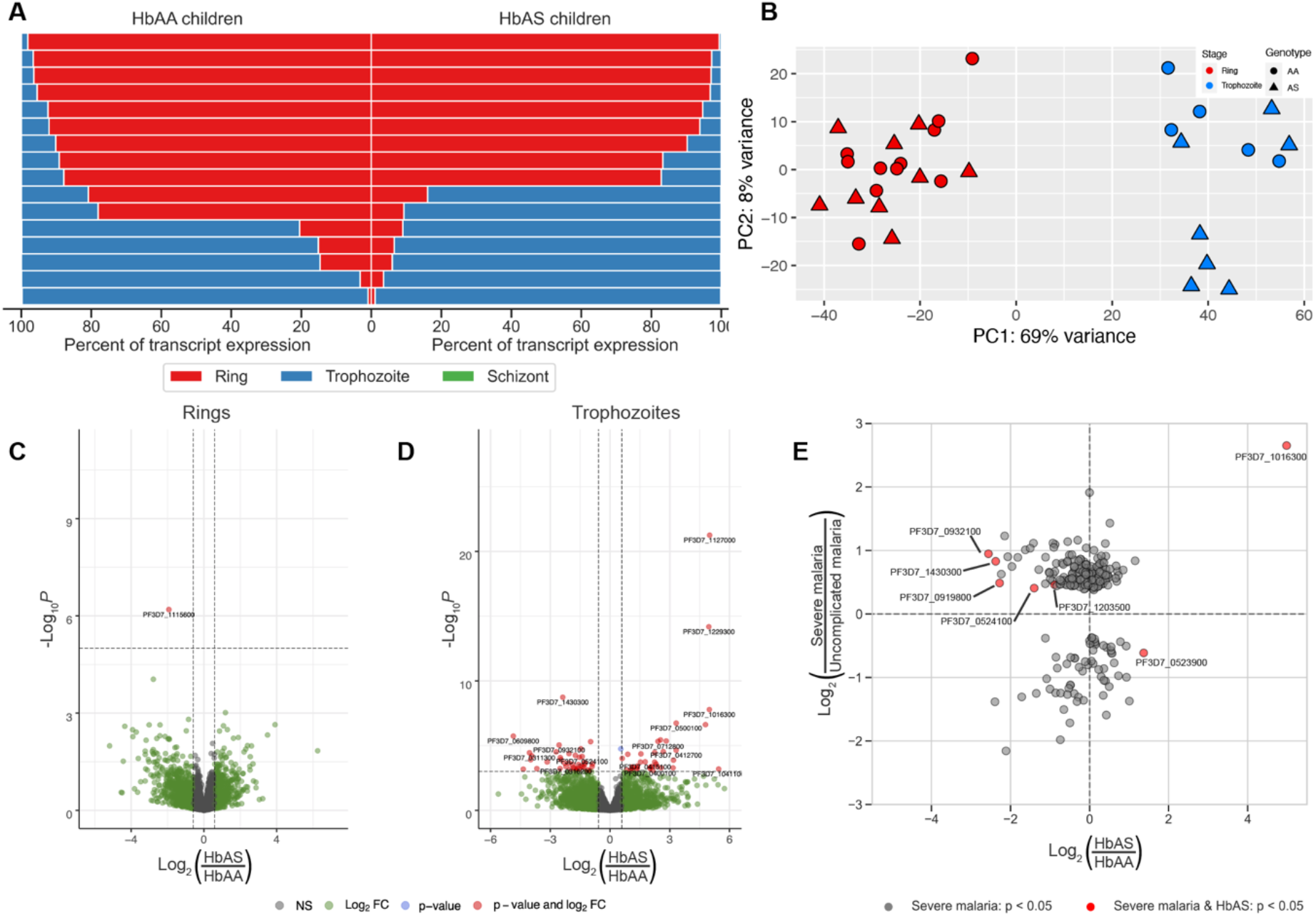
Differential expression profiling of parasite transcripts in freshly-collected infections in Malian children with uncomplicated malaria with HbAA or HbAS. **(A)** Proportion of parasite stage-specific transcripts expressed in each patient’s infection (rows) in HbAA (left) and HbAS (right) children. Expression of stage-restricted transcripts for ring (red), trophozoite (blue), and schizont (green) stages were measured as TPMs, summed within each stage, and expressed as a proportion of stage for each infection. We observed a near complete absence of schizont-specific transcript expression, and most infections were predominantly either rings or trophozoites with little admixture. See Jupyter Notebook “Genes_by_stage.ipynb” for full workflow. **(B)** Principal component analysis of transcript expression from *in vivo* isolates following variance-stabilizing transformation based on the 500 transcripts with the highest variance across all samples. Clusters separate according to parasite stage identified from stage-specific transcript expression. **(C)** Volcano plots of differential transcript expression conditioned by parasite stage. *X*-axis is the log2 fold change of HbAS versus HbAA children, with positive values indicating increased expression in HbAS. *Y*-axis is −log10 p-values of each transcript, and horizontal dashed lines indicate p-value thresholds that capture the transcripts with p-adjusted values < 0.05. **(E)** Scatterplots of the log_2_ fold-change of individual transcripts for severe malaria parasites compared to uncomplicated malaria parasites in Gambian children (*y*-axes) and HbAS compared to HbAA parasites (*x*-axes). Points in the upper right and lower left quadrants signify transcripts that are differentially expressed in the same direction between studies, i.e. those in the upper right quadrant are upregulated in both severe malaria and HbAS, and those in the lower left quadrant are downregulated in both severe malaria and HbAS. All transcripts included were statistically significant in severe malaria versus uncomplicated malaria from (27). Red points have an adjusted p-value < 0.05 in trophozoite parasites from HbAS vs HbAA Malian children.

### Differential expression *in vivo*

The only differentially-expressed transcript in ring-stage infections was cyclophilin 19B (*cyp19B*, PF3D7_1115600), a peptidyl-prolyl cis-trans isomerase (PPIase) (**Figure 6C, Supplementary Table 2**): compared to children with HbAA, *cyp19B* in those with HbAS was downregulated threefold (p-adjusted = 0.023010). qPCR confirms that *cyp19B* is reduced in ring-stage parasite transcriptomes in HbAS (**Figure S15**). Cyclophilins function in protein folding and trafficking, RNA processing, and RNA-induced silencing complex (RISC) assembly (22) and is most highly expressed in schizont-stage parasites *in vitro* (23).

We observed in trophozoite-stage patient samples differential expression of 74 transcripts (33 upregulated, 41, downregulated) (**Figure 6D, Supplementary Table 2**). We did not observe any enrichment for GO terms amongst the upregulated or downregulated transcripts. The most upregulated transcript in HbAS trophozoites was a putative protein phosphatase (PF3D7_1127000, log2 fold change = 5.00, p-adjusted = 1.08×10^−18^). Also among the 10 most upregulated transcripts in HbAS trophozoites were the 130 kD glycophorin binding protein (GBP130, PF3D7_1016300) and the two glycophorin binding protein homologs (GBPH, PF3D7_1401000 and GBPH2, PF3D7_1301200), each upregulated greater than eight-fold in HbAS trophozoites. Additionally, one of the most significantly upregulated transcripts in HbAS trophozoites (log2 fold change = 1.55, p-adjusted = 0.0077) encodes the heat shock protein 70-x (HSP70-x, PF3D7_08317000), which is thought to chaperone export of PfEMP-1 via J-dots and Maurer’s clefts (24), similar to the 2-fold overexpression of the critical Maurer’s cleft components MAHRP1 (PF3D7_1370300, p-adjusted = 0.041) and MAHRP2 (PF3D7_1353200, p-adjusted = 0.034).

Among the 41 downregulated transcripts in trophozoites in HbAS children, the most highly significant was that encoding a putative acid phosphatase (PF3D7_1430300, log2 fold change = −2.37, p-adjusted = 1.77×10^−6^). We also observed among the top 10 downregulated transcripts the putative autophagy protein 5 (ATG5) (PF3D7_1430400, p-adjusted = 0.0076) which associates with autophagosome-like structures,(25) and the DNA repair protein RAD5 (PF3D7_1343400, p-adjusted = 0.0091), which has been linked to delayed clearance following artemisinin treatment (26).

Because HbAS confers protection specifically from severe malaria, we compared our parasite gene expression in Malian children with that of a prior study comparing parasite gene expression in Gambian children with severe and uncomplicated malaria (27), which identified 236 differentially-expressed transcripts. Seven of these transcripts were also differentially expressed between trophozoite-stage parasites in HbAS and HbAA children (**Figure 6E**). Of these seven, six were transcripts overexpressed in severe malaria. One of these, *gbp130* (PF3D7_1016300), was upregulated in children with HbAS. Five of the six transcripts that were upregulated in severe malaria were downregulated in children with HbAS (PF3D7_0932100, PF3D7_1203500, PF3D7_0524100, PF3D7_0919800, PF3D7_1430300). Among the transcripts with statistically-significant upregulation in severe malaria but downregulation in HbAS RBCs was that of a putative acid phosphatase (PF3D7_1430300), which was the most statistically significant downregulated transcript in our HbAS trophozoite samples compared to HbAA trophozoites.

### Intersection of aberrant transcripts in HbAS across analytic approaches

Across analyses of both differential expression and temporal dysregulation in our time-series experiments, only 15 candidate transcripts were aberrantly expressed in HbAS by 3D7 and FUP parasites (**Figure 7A**). These include PTEX88, which was upregulated in both 3D7 and FUP, which is noteworthy owing to its critical role in exporting proteins from the parasitophorous vacuole membrane (PVM) that remodel the iRBC. Of these candidates, 10 overlapped with candidates identified from *in vivo* infections in the same direction (upregulated/downregulated, **Supplementary Table 2**). The sole transcript differentially expressed in *in vivo* ring-stage parasites, *cyp19B*, was also downregulated in both 3D7 and FUP HbAS (**Figure 7B, 7C, 7D**).

**Fig 7.**
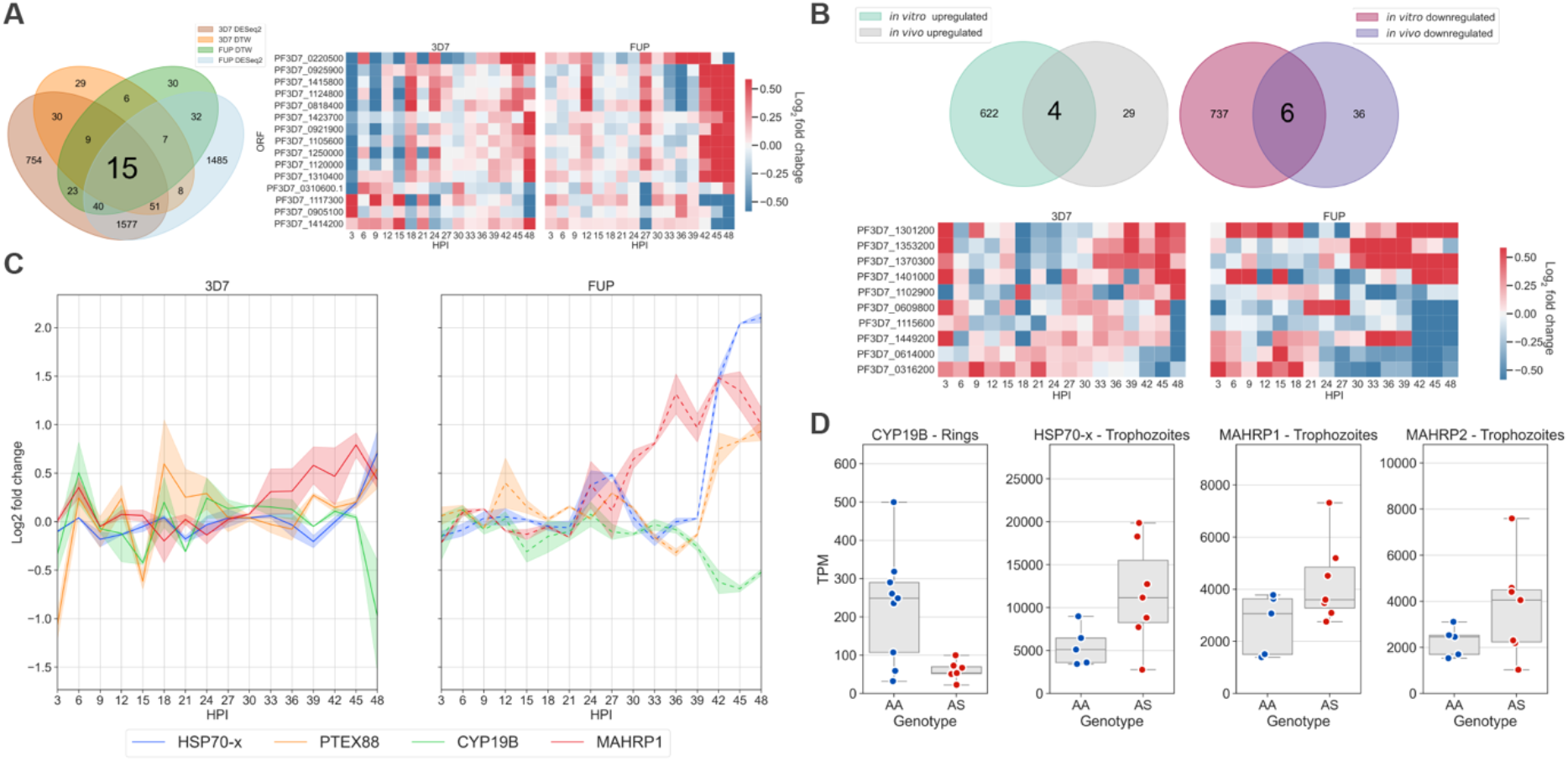
Intersection of aberrant transcripts across analytical and experimental approaches. **(A)** Venn diagram of transcripts that were differentially expressed and temporally dysregulated in 3D7 and FUP in *in vitro* time series experiments. The 15 transcripts that were differentially expressed and temporally dysregulated in 3D7 and FUP are visualized in a heatmap of log2 fold changes between HbAS:HbAA samples over 48 hours. This list includes seven unannotated transcripts (PF3D7_0921900, PF3D7_1423700, PF3D7_1117300, PF3D7_1414200, PF3D7_0925900, PF3D7_1120000, PF3D7_1310400), an exported protein (PF3D7_0220500), two rRNA processing proteins (PF3D7_1250000, PF3D7_0818400), a translation initiation factor (PF3D7_0310600.1), a nuclear preribosomal assembly protein (PF3D7_1124800), a nucleoporin (PF3D7_0905100), a dimethyladenosine transferase (PF3D7_1415800), and the translocon component PTEX88 (PF3D7_1105600). **(B)** Venn diagram of transcripts that are up- and downregulated in both *in vitro* time series and *in vivo* samples from Malian children. The 10 transcripts that are shared between both experiments are visualized in a heatmap, constrained to the same conditions specified in **(A). (C)** Log2 fold change of transcripts over the *in vitro* time series of transcripts of biological significance that were included in **(A)** and **(B). (D)** Normalized expression of transcripts of transcripts of biological significance in *in vivo* samples from **(B)**.

Among the genes that were significantly upregulated *in vitro* and *in vivo* was *gbp130*, whose expression was increased more than 30-fold in HbAS trophozoites compared to HbAA. In our *in vitro* time series, we find this gene is significantly upregulated in HbAS FUP at 48 hpi, and one HbAS 3D7 replicate also demonstrated upregulation at 48 hpi. In addition, the glycophorin binding protein homologs, *gbph* and *gbph2*, were significantly upregulated *in vivo* in trophozoite-stage parasite isolates and *in vitro* in both HbAS 3D7 and HbAS FUP. *In vitro, hsp70-x* (PF3D7_0831700) demonstrated a pattern of elevated transcript abundance at late time points in 3D7 and FUP, although statistical significance was only achieved in FUP (**Figure 7C, 7D**). Additionally, *mahrp1* and *mahrp2* were upregulated in HbAS *in vitro* and *in vivo* (**Figure 7C, 7D**). Among downregulated transcripts *in vivo* and *in vitro*, we observed a palmitoyltransferase (PF3D7_0609800), a thiamin-phosphate pyrophosphorylase (PF3D7_0614000), an exported protein (hyp11, PF3D7_1102900) and two unannotated proteins of unknown function (PF3D7_0316200, PF3D7_1449200). The protein phosphatase (PF3D7_1127000) that was the most highly upregulated transcript in HbAS trophozoites was highly expressed in HbAS, and also found highly expressed and slightly upregulated in HbAS FUP, although it did not pass the threshold for statistical significance. In 3D7, however, this transcript was lowly expressed in all samples. Finally, we examined the set of candidate aberrant transcripts identified *in vivo* that were similarly expressed between Hb genotypes *in vitro*, a set which may indicate parasite responses that are specific to physiologic conditions. These 17 *in vivo-*specific transcripts include upregulation of *atg5* (PF3D7_1430400) and *rad5* (PF3D7_PF3D7_1343400), suggesting autophagy and DNA repair, as well as the differential expression of enzymes whose specific functions are currently not well understood, including the putative protein phosphatase PF3D7_112700, and the putative acid phosphatase PF3D7_1430300, are impacted by HbAS *in vivo*.

## Discussion

Across studies, HbAS consistently protects against severe *P. falciparum* malaria by mechanisms that remain obscure. In this study, we investigated the transcriptional changes that occur in *P. falciparum* in response to HbAS. Overall, we found that the parasite’s transcriptional program remains largely unperturbed in HbAS iRBCs. However, hundreds of transcripts demonstrated differential expression in HbAS, with the bulk of these changes occurring late in the IDC. Combining our *in vitro* experiments with parasite transcript expression *in vivo,* we observed impacts of HbAS on transcripts encoding the parasite’s protein chaperone and folding machinery, oxidative stress response, and protein export machinery.

Transcripts that were most clearly temporally dysregulated in HbAS iRBCs were those that are components of gene regulation: the GO biological processes with the highest DTW scores were those for transcription by RNA polymerase II, RNA splicing, regulation of translation initiation, and ribosome biogenesis. Mis-timed expression of these transcripts could result in a cascade of downstream expression changes. In particular, temporal dysregulation of the subunits of RNA polymerase II, the complex responsible for the production of all parasite mRNA, could result in altered transcript abundance. Furthermore, the downstream impact of temporal dysregulation of transcripts involved in RNA splicing, ribosome biogenesis, and translation initiation could lead to significant alterations in the HbAS parasite proteome.

Differential expression in HbAS emerged primarily during the schizont stage in our *in vitro* time series. The molecular functions and cellular components that were most downregulated in HbAS include cysteine-type peptidase activity (in particular the transcripts encoding the SERA proteins), and the core histones and histone variants (**Figure 3C**). The SERA proteins are thought to play a role in erythrocyte rupture due to their localization to the parasitophorous vacuole (PV) in late stages of the IDC and to a papain-like cysteine peptidase domain that is common to all SERA proteins (28). The highly-expressed *SERA5* (29) along with *SERA6* are essential to parasite growth in the blood stage of infection (30, 31), and conditional inactivation of *SERA5* disrupts schizont rupture and merozoite release (32). Disruption of individual *SERA* expression can result in compensatory overexpression of other SERAs (33), though our observation of consistent and significant downregulation of *SERA2*, *−3*, *−4*, *−5*, *−6*, and *−7* (**Figure 4C**) precludes this in HbAS and suggests that SERA-mediated egress is impaired in HbAS. In addition to efficient egress, IDC propagation relies on a highly regulated epigenome (34), which is normally regulated by histones transcribed late in the asexual blood stage of development for nucleosome packaging in merozoites (35, 36). We observed that transcripts encoding *P. falciparum* histones are consistently under-expressed in HbAS iRBCs (**Figure 4B**), which could produce an altered chromatin landscape in the new merozoites and alter the blood stage transcriptional program after subsequent invasions. Though we did not observe large transcriptional changes in *in vivo* parasites from HbAS children, such an effect *in vivo* could be masked if such a defective transcriptional program inhibits these merozoites from successfully parasitizing an RBC. Taken together, these observations lead to the possibility that HbAS impairs the egress of new merozoites mediated by SERA5, and that a subset of merozoites are loaded with aberrantly packaged nucleosomes, collectively reducing the efficiency of propagation. This could explain why HbAS is associated with lower parasite densities *in vivo* (37–39). ß-globin variants, including HbAS, result in dysmorphic and dysfunctional Maurer’s clefts (8, 40, 41). Surprisingly, we found that transcripts related to Maurer’s clefts were among the most upregulated transcripts both *in vitro* and *in vivo*. *P. falciparum* exports a large number of proteins, including PfEMP-1, to the iRBC surface in knob-like protrusions that mediate its cytoadherence. Maurer’s clefts serve as the hub through which these proteins are trafficked (42). In particular, *sbp1*, *mahrp1*, and *mahrp2* were upregulated in both of our *in vitro* strains, with the latter two also upregulated *in vivo* in trophozoite stage parasites (**Figure 7D**). In the asexual blood stage, *sbp1* and *mahrp1* are non-essential, but individual disruption of either gene leads to dysmorphic Maurer’s clefts and the absence of PfEMP-1 on the iRBC surface (43, 44). In the IDC, *mahrp2* is essential, and the MAHRP2 protein localizes to electron-dense tubular structures known as tethers that connect Maurer’s clefts to the parasitophorous vacuole membrane (PVM) and the RBC membrane (45, 46). These data suggest the possibility that gene expression of Maurer’s clefts is under transcription-translation feedback control that is contingent upon the proper downstream assembly and function of its protein complexes. In this scenario, parasites in HbAS iRBCs would continue to express Maurer’s cleft genes to compensate for the dysmorphic and dysfunctional Maurer’s clefts observed in this context.

Of particular interest among the six transcripts that were significantly downregulated in HbAS both *in vitro* and *in vivo* was *cyp19B*, which in our *in vivo* samples was approximately 3-fold downregulated. CYP19B is found primarily in the parasite cytosol and exhibits PPIase activity, participating in protein folding and potentially acting as a molecular chaperone via protein trafficking and regulation of multi-protein complexes (23). Like several other cyclophilins, CYP19B is bound by Cyclosporin A (CsA), which inhibits *in vitro* both the PPIase activity of CYP19B (47) as well as the growth of *P. falciparum* (23). Interestingly, *cyp19B* was identified as the most upregulated gene in Artemisinin-resistant (Art-R) parasites in a Southeast Asian population transcriptomic comparison of over 1000 *in vivo P. falciparum* isolates, in which its expression was the most positively correlated single transcript with increased parasite clearance half-life (48). In that context, its upregulation correlated with a robust parasite stress response to the free-radical damage initiated by artemisinin. In contrast, in our experiments *cyp19B* was under-expressed in HbAS RBCs, despite the presumptively oxidizing and harsh environment of the HbAS RBC. CYP19B is also a member of the Plasmodium reactive oxidative stress complex (PROSC), and we observed a downregulation of a majority of the members of PROSC in HbAS *in vitro* at late time points in the IDC in both 3D7 and FUP, including *cyp19B*, *hsp70-2*, *grp94*, *pdi8*, and *pferc* (PF3D7_1115600, PF3D7_0917900, PF3D7_1222300, PF3D7_0827900, PF3D7_1108600), though not i*n vivo.* Additionally, the relatively high levels of *cyp19B* expression in HbAA *in vivo* ring-stage samples was unexpected, given that both transcript expression (in our *in vitro* time series) and protein expression (49) are highest in the late trophozoite and schizont stages.

Among our *in vivo* parasites, parasites were either predominantly rings or trophozoites with no minor population > 20%, suggesting that circulating parasites in *in vivo* infections are broadly synchronized. This divergence accounted for most of the variance in transcriptomes, underscoring the need for stage-matched comparisons of parasite transcription. *In vitro*, most of the transcriptional changes we observed occurred in schizont-stage parasites. However, given that schizont-stage parasites are likely sequestered in the microvasculature and thus not in circulation, we could not confirm that similar changes occur *in vivo*. This does not mean, though, that parasites in HbAS iRBCs *in vivo* would not be similarly impacted. Finally, though we observed relatively few differentially-expressed transcripts in HbAS children – particularly among the ring stages that predominated – the intersection of statistically-significant differential expression between our trophozoite-stage data compared with transcriptomic data from severe malaria indicates that HbAS reduces the expression of a subset of transcripts that are associated with severe malaria. The functions of these transcripts are largely obscure but our analyses collectively suggest their functional role in severe malaria. Despite the expectation that HbAS would reduce microvascular cytoadherence and thereby produce more circulating trophozoites, we observed similar proportions of trophozoite-predominant infections in HbAS (7/16) and HbAA children (5/16) (**Fig. 6A, 6B**), and no evidence of late trophozoite or schizont gene expression in our HbAS patient samples (**Fig 6A, S13**). This may have been influenced by our sampling scheme, in which samples were collected the day following a febrile episode, and since parasites are thought to be somewhat synchronous in natural infections (50) and our data suggest, we would expect to find only ring-stage and early trophozoite parasites in our patients.

It is surprising that we observed conservation of the overall order of the transcriptional program in the IDC despite the presence of substantial temporal dysregulation in HbAS of gene regulatory GO terms such as RNA polymerase II and translation initiation. Interestingly, our *in vitro* time series data in HbAS RBCs mirrored key aspects of the transcriptional changes observed in *P. falciparum* lines mutated for drug-resistance transporter genes *pfmdr-1* and *pfcrt* (51), in which parasites demonstrated an overall concordance in the temporal expression of the IDC transcriptome, with most of the transcriptional changes impacting the abundance of late-stage transcripts. This suggests the possibility that *P. falciparum’s* transcriptional program in the IDC is sufficiently robust to adapt to fitness challenges posed by HbAS RBCs and aspects related to drug resistance; these adaptations primarily manifest by modulating the expression levels of particular genes, while still producing the necessary transcripts on time to complete the IDC. The maintenance of the parasite’s transcriptional program relies on a robust architecture that is only partially understood, highlighted by recent reports that *Plasmodium spp.* possess intrinsic oscillators that govern the periodicity of their IDC program (52, 53). The alterations we observed in transcript abundances may be a consequence of the temporal dysregulation of gene regulatory products we discovered with DTW analysis. Such subtle alterations could, with successive rounds of infection, be amplified into fitness-mediating phenotypic changes.

Our study had several limitations. Our *in vitro* time series experiments were conducted for one round of infection over 48 hours in order to ensure tight synchrony of parasites, and therefore we cannot observe impacts of perturbations in histones and chromatin modifications on subsequent infections. Our *in vitro* experiments were performed in single gas mixture at 1% [O_2_], while *in vivo*, *P. falciparum* encounters in circulation a range of oxygen tension that can differentially impact the growth of *P. falciparum* in HbAS iRBCs *in vitro* (3–6). Though prior studies have reported a growth defect in HbAS RBCs at 1% [O_2_], we did not observe such a defect over one IDC.

Our data encompass *P. falciparum’s* transcriptional response to infection in HbAS RBCs, highlighting multiple avenues by which HbAS neutralizes *P. falciparum*. Our survey of parasite transcriptomes reveals that key components of Maurer’s clefts are significantly upregulated in HbAS, which could signal a compensatory response to the dysmorphic and disorganized Maurer’s clefts that lead to altered surface knobs and diminished cytoadherence previously identified in HbAS. In addition, the significant reduction of SERA and nucleosome transcripts in HbAS indicate the possibility that daughter merozoites are inhibited in efficient egress and inherit aberrantly packaged DNA in histones. Furthermore, downregulation of *cyp19B* in HbAS potentially makes the parasite more susceptible to the stressful environment of an HbAS RBC. These transcriptional perturbations induced in HbAS provide new targets to neutralize parasite mediators of disease, and suggest that downstream changes in the proteome and epigenome are candidates for exploration.

## Materials and Methods

### Subjects and samples

HbAA and HbAS blood samples were collected at Duke University Hospital under IRB # Pro00007816 for *in vitro* experiments. HbAS genotypes were confirmed by Sanger sequencing. Field samples were collected in an observational study of the efficacy of Artemether-Lumefantrine for the treatment of uncomplicated malaria in children in Kéniéroba, Mali (https://ClinicalTrials.gov Identifier: NCT02645604), where HbAS reduces the risk of uncomplicated malaria by 34% (18). Inclusion criteria for this study was age between 2 and 17 years of age and febrile uncomplicated falciparum malaria as assessed by light microscopy. From all participants prior to treatment, we collected venous blood, passed this through cellulose columns (54), and stored up to 2mL of the flow-through in RNAprotect (Qiagen). Parasite density was estimated from light microscopy, hemoglobin genotype assessed using HPLC, and ABO blood group by agglutination assay. For transcriptional analysis, we selected all available samples from children with HbAS and matched each of these to a sample from a child with HbAA on: month of episode, parasite density, ethnic background, and, if possible, ABO blood type. The field study was approved by the Institutional Review Board of the University of Sciences, Techniques, and Technologies of Bamako (IRB00001983).

### Parasite culture

*P. falciparum* parasites of 3D7 and FUP strains were cultured in human RBCs in RPMI-1640 supplemented with Albumax II, L-Glutamine, Hypoxanthine, and D-Glucose at a 2% hematocrit in 1% O_2_, 5% CO_2_ in N_2_ at 37°C (55). iRBCs were kept in filtered-lid cell culture flasks inside a modular hypoxia incubator chamber and mixed on a nutator. *P. falciparum* parasites from either strain were synchronized using Percoll centrifugation, and the schizont fraction was then split equally into separate flasks of HbAS or HbAA. Following a 3-hour incubation, these cultures were treated with Sorbitol, and allowed to progress through the intraerythrocytic cycle in parallel, with sampling every 3 hours for light microscopy and RNA preservation. Parasite density and maturation was assessed following Giemsa staining by readers masked to hemoglobin type and hours post invasion (hpi).

### Library preparation

Total RNA was isolated using TRIzol and the RNeasy Mini Kit (Qiagen) after DNase treatment, and quantified by Qubit High Sensitivity RNA Assay (Thermo Fisher Scientific) before storage at −80°C. Libraries were prepared from total RNA with the Kapa Stranded mRNA-seq library prep kit. Libraries from each of the two time series experiments (n=64/experiment) were sequenced on a full flow cell on the NovaSeq 6000 S2 platform with 150 base pair (bp) paired-end reads. Field study samples (n=32) were sequenced on a full flow cell of the NovaSeq 6000 S1 platform with 50bp paired-end reads.

### Transcriptome quantification

Reads were assessed for quality with FastQC, trimmed and quality-filtered with Trimmomatic (56), depleted of reads mapping to the human genome (GRCh38.p13) with STAR (57), and mapped and quantified with Salmon (58) using the *Plasmodium falciparum* 3D7 transcriptome (ASM276v2). Quantification files of parasite gene expression were summarized with the *tximport* package in R (59). Principal component analysis data was generated in R with the plotPCA function from DESeq2 and then plotted with ggplot2. For heatmap visualization, transcripts per million (TPMs) were normalized around the mean for each transcript over the time series and depicted as a *z* score of standard deviations from the mean.

### Peak shift analysis

Analyses of the shift of transcript peak were conducted by first identifying for each transcript in each time series the timepoint at which the transcript achieved its maximum expression, and then computing for each transcript the difference between each time series of the transcript’s peak. We repeated this approach using only transcripts with a single peak over the time series, which were identified using the find_peaks function from the signal processing submodule in SciPy. We compared the distributions of peak shift values between samples using Kruskal-Wallis one-way analysis of variance.

### Differential expression analysis

All differential expression analyses were performed with the DESeq2 (60) R package using unnormalized read counts per transcript. Transcripts were considered differentially expressed if, as determined by DESeq2 with a false-discovery rate set to 0.05, both their adjusted p-value < 0.05 and |log_2_(HbAS/HbAA)| > log_2_(1.5) in the time-series, and with a false-discovery rate set to 0.1, both the adjusted p-value < 0.05 and |log_2_(HbAS/HbAA)| > log_2_(1.5) in the field samples.

### Gene ontology enrichment analysis

Gene ontology enrichment analysis was performed with GOATOOLS (17). Gene IDs and associated GO IDs were downloaded from NCBI (ftp://ftp.ncbi.nlm.nih.gov/gene/DATA/gene2go.gz), and *P. falciparum’s* annotations were extracted with the taxonomic identifier 36329. Using the transcripts identified as differentially expressed by DESeq2 according to the conditions described above as input, gene ontology terms were accepted for Benjamini-Hochberg adjusted p-values < 0.05.

### Analyses of transcript temporal regulation

Dynamic time warping (DTW) of time series transcript expression data was computed with the tslearn toolkit (61). We considered as temporally-dysregulated transcripts with mean DTW scores across ß-globin replicates that exceeded two standard deviations above the mean DTW scores of all transcripts in HbAA versus HbAS comparisons. We used the Temporal Alignment Kendall-Tau algorithm (TAKT) (19) to identify transcripts within each parasite that had similar dynamics but were temporally shifted between β-globin types during ring stage (19). Measures of similarity and associated significance scores were produced for a gene between similar β-globin type replicates and a p-value of ≤ 0.05 was used to identify transcripts with similar dynamics across all β-globin types within each parasite.

### Stage classification of *in vivo* parasites

We identified the set of transcripts shared between 3D7 and FUP from our *in vitro* time series that peaked within rings, trophozoites, or schizonts, and then selected transcripts that were highly expressed at their peak, discarding those that did not exceed 1,000 TPM. We also estimated parasite stage with a previously published mixture model (21). Our count data was converted to RPKM and log2-transformed to match the staged expression data (62) used for stage estimation.

## Acknowledgments

We thank Marilyn Telen and Nirmish Shah (both of Duke University) for their assistance with blood collection, and Tom Wellems and Chanaki Amaratunga (both of NIAID) for assistance with clinical study data. *P. falciparum* strains 3D7 (MRA-102, contributed by Daniel J. Carucci) and FUP UGANDA-PALO ALTO (MRA-915, contributed by T. Sam-Yellowe) were obtained from BEI Resources, NIAID, NIH. We are indebted to the Malian children and their guardians who participated in the field study, and to our donors of normal and sickle-trait blood.

## References

1. Malaria Genomic Epidemiology N, Malaria Genomic Epidemiology N. Reappraisal of known malaria resistance loci in a large multicenter study. Nat Genet. 2014;46(11):1197–204.

2. Taylor SM, Parobek CM, Fairhurst RM. Haemoglobinopathies and the clinical epidemiology of malaria: a systematic review and meta-analysis. The Lancet infectious diseases. 2012;12(6):457–68.

3. Pasvol G. The interaction between sickle haemoglobin and the malarial parasite Plasmodium falciparum. Transactions of the Royal Society of Tropical Medicine and Hygiene. 1980;74(6):701–5.

4. Friedman MJ. Erythrocytic mechanism of sickle cell resistance to malaria. Proceedings of the National Academy of Sciences of the United States of America. 1978;75(4):1994–7.

5. Pasvol G, Weatherall DJ, Wilson RJ. Cellular mechanism for the protective effect of haemoglobin S against P. falciparum malaria. Nature. 1978;274(5672):701–3.

6. Archer NM, Petersen N, Clark MA, Buckee CO, Childs LM, Duraisingh MT. Resistance to Plasmodium falciparum in sickle cell trait erythrocytes is driven by oxygen-dependent growth inhibition. Proceedings of the National Academy of Sciences of the United States of America. 2018;115(28):7350–5.

7. LaMonte G, Philip N, Reardon J, Lacsina JR, Majoros W, Chapman L, et al. Translocation of sickle cell erythrocyte microRNAs into Plasmodium falciparum inhibits parasite translation and contributes to malaria resistance. Cell host & microbe. 2012;12(2):187–99.

8. Cyrklaff M, Srismith S, Nyboer B, Burda K, Hoffmann A, Lasitschka F, et al. Oxidative insult can induce malaria-protective trait of sickle and fetal erythrocytes. Nature communications. 2016;7:13401.

9. Waldecker M, Dasanna AK, Lansche C, Linke M, Srismith S, Cyrklaff M, et al. Differential time-dependent volumetric and surface area changes and delayed induction of new permeation pathways in P. falciparum-infected hemoglobinopathic erythrocytes. Cellular microbiology. 2017;19(2).

10. Cholera R, Brittain NJ, Gillrie MR, Lopera-Mesa TM, Diakite SA, Arie T, et al. Impaired cytoadherence of Plasmodium falciparum-infected erythrocytes containing sickle hemoglobin. Proceedings of the National Academy of Sciences of the United States of America. 2008;105(3):991–6.

11. Kilian N, Srismith S, Dittmer M, Ouermi D, Bisseye C, Simpore J, et al. Hemoglobin S and C affect protein export in Plasmodium falciparum-infected erythrocytes. Biol Open. 2015;4(3):400–10.

12. Lansche C, Dasanna AK, Quadt K, Frohlich B, Missirlis D, Tetard M, et al. The sickle cell trait affects contact dynamics and endothelial cell activation in Plasmodium falciparum-infected erythrocytes. Commun Biol. 2018;1:211.

13. Fairhurst RM, Bess CD, Krause MA. Abnormal PfEMP1/knob display on Plasmodium falciparum-infected erythrocytes containing hemoglobin variants: fresh insights into malaria pathogenesis and protection. Microbes Infect. 2012;14(10):851–62.

14. Serjeant GR. The natural history of sickle cell disease. Cold Spring Harb Perspect Med. 2013;3(10):a011783.

15. Preston MD, Campino S, Assefa SA, Echeverry DF, Ocholla H, Amambua-Ngwa A, et al. A barcode of organellar genome polymorphisms identifies the geographic origin of Plasmodium falciparum strains. Nat Commun. 2014;5:4052.

16. Geiman QM, Meagher MJ. Susceptibility of a New World monkey to Plasmodium falciparum from man. Nature. 1967;215(5099):437–9.

17. Klopfenstein DV, Zhang L, Pedersen BS, Ramirez F, Warwick Vesztrocy A, Naldi A, et al. GOATOOLS: A Python library for Gene Ontology analyses. Sci Rep. 2018;8(1):10872.

18. Lopera-Mesa TM, Doumbia S, Konate D, Anderson JM, Doumbouya M, Keita AS, et al. Effect of red blood cell variants on childhood malaria in Mali: a prospective cohort study. Lancet Haematol. 2015;2(4):e140–9.

19. Kelliher CM, Foster MW, Motta FC, Deckard A, Soderblom EJ, Moseley MA, et al. Layers of regulation of cell-cycle gene expression in the budding yeast Saccharomyces cerevisiae. Mol Biol Cell. 2018;29(22):2644–55.

20. Miller LH, Baruch DI, Marsh K, Doumbo OK. The pathogenic basis of malaria. Nature. 2002;415(6872):673–9.

21. Tonkin-Hill GQ, Trianty L, Noviyanti R, Nguyen HHT, Sebayang BF, Lampah DA, et al. The Plasmodium falciparum transcriptome in severe malaria reveals altered expression of genes involved in important processes including surface antigen-encoding var genes. PLoS Biol. 2018;16(3):e2004328.

22. Kumari S, Roy S, Singh P, Singla-Pareek SL, Pareek A. Cyclophilins: proteins in search of function. Plant Signal Behav. 2013;8(1):e22734.

23. Gavigan CS, Kiely SP, Hirtzlin J, Bell A. Cyclosporin-binding proteins of Plasmodium falciparum. Int J Parasitol. 2003;33(9):987–96.

24. Kulzer S, Charnaud S, Dagan T, Riedel J, Mandal P, Pesce ER, et al. Plasmodium falciparum-encoded exported hsp70/hsp40 chaperone/co-chaperone complexes within the host erythrocyte. Cell Microbiol. 2012;14(11):1784–95.

25. Joy S, Thirunavukkarasu L, Agrawal P, Singh A, Sagar BKC, Manjithaya R, et al. Basal and starvation-induced autophagy mediates parasite survival during intraerythrocytic stages of Plasmodium falciparum. Cell Death Discov. 2018;4:43.

26. Takala-Harrison S, Clark TG, Jacob CG, Cummings MP, Miotto O, Dondorp AM, et al. Genetic loci associated with delayed clearance of Plasmodium falciparum following artemisinin treatment in Southeast Asia. Proceedings of the National Academy of Sciences of the United States of America. 2013;110(1):240–5.

27. Lee HJ, Georgiadou A, Walther M, Nwakanma D, Stewart LB, Levin M, et al. Integrated pathogen load and dual transcriptome analysis of systemic host-pathogen interactions in severe malaria. Sci Transl Med. 2018;10(447).

28. Rosenthal PJ. Cysteine proteases of malaria parasites. International journal for parasitology. 2004;34(13-14):1489–99.

29. Aoki S, Li J, Itagaki S, Okech BA, Egwang TG, Matsuoka H, et al. Serine repeat antigen (SERA5) is predominantly expressed among the SERA multigene family of Plasmodium falciparum, and the acquired antibody titers correlate with serum inhibition of the parasite growth. J Biol Chem. 2002;277(49):47533–40.

30. Miller SK, Good RT, Drew DR, Delorenzi M, Sanders PR, Hodder AN, et al. A subset of Plasmodium falciparum SERA genes are expressed and appear to play an important role in the erythrocytic cycle. J Biol Chem. 2002;277(49):47524–32.

31. Thomas JA, Collins CR, Das S, Hackett F, Graindorge A, Bell D, et al. Development and Application of a Simple Plaque Assay for the Human Malaria Parasite Plasmodium falciparum. PLoS One. 2016;11(6):e0157873.

32. Collins CR, Hackett F, Atid J, Tan MSY, Blackman MJ. The Plasmodium falciparum pseudoprotease SERA5 regulates the kinetics and efficiency of malaria parasite egress from host erythrocytes. PLoS Pathog. 2017;13(7):e1006453.

33. McCoubrie JE, Miller SK, Sargeant T, Good RT, Hodder AN, Speed TP, et al. Evidence for a common role for the serine-type Plasmodium falciparum serine repeat antigen proteases: implications for vaccine and drug design. Infect Immun. 2007;75(12):5565–74.

34. Chaal BK, Gupta AP, Wastuwidyaningtyas BD, Luah YH, Bozdech Z. Histone deacetylases play a major role in the transcriptional regulation of the Plasmodium falciparum life cycle. PLoS pathogens. 2010;6(1):e1000737.

35. Coetzee N, Sidoli S, van Biljon R, Painter H, Llinas M, Garcia BA, et al. Quantitative chromatin proteomics reveals a dynamic histone post-translational modification landscape that defines asexual and sexual Plasmodium falciparum parasites. Sci Rep. 2017;7(1):607.

36. Ponts N, Harris EY, Prudhomme J, Wick I, Eckhardt-Ludka C, Hicks GR, et al. Nucleosome landscape and control of transcription in the human malaria parasite. Genome Res. 2010;20(2):228–38.

37. Aidoo M, Terlouw DJ, Kolczak MS, McElroy PD, ter Kuile FO, Kariuki S, et al. Protective effects of the sickle cell gene against malaria morbidity and mortality. Lancet. 2002;359(9314):1311–2.

38. Le Hesran JY, Personne I, Personne P, Fievet N, Dubois B, Beyeme M, et al. Longitudinal study of Plasmodium falciparum infection and immune responses in infants with or without the sickle cell trait. International journal of epidemiology. 1999;28(4):793–8.

39. Stirnadel HA, Stockle M, Felger I, Smith T, Tanner M, Beck HP. Malaria infection and morbidity in infants in relation to genetic polymorphisms in Tanzania. Trop Med Int Health. 1999;4(3):187–93.

40. Cyrklaff M, Sanchez CP, Kilian N, Bisseye C, Simpore J, Frischknecht F, et al. Hemoglobins S and C interfere with actin remodeling in Plasmodium falciparum-infected erythrocytes. Science. 2011;334(6060):1283–6.

41. Kilian N, Dittmer M, Cyrklaff M, Ouermi D, Bisseye C, Simpore J, et al. Haemoglobin S and C affect the motion of Maurer’s clefts in Plasmodium falciparum-infected erythrocytes. Cellular microbiology. 2013;15(7):1111–26.

42. Lanzer M, Wickert H, Krohne G, Vincensini L, Braun Breton C. Maurer’s clefts: a novel multi-functional organelle in the cytoplasm of Plasmodium falciparum-infected erythrocytes. Int J Parasitol. 2006;36(1):23–36.

43. Spycher C, Rug M, Pachlatko E, Hanssen E, Ferguson D, Cowman AF, et al. The Maurer’s cleft protein MAHRP1 is essential for trafficking of PfEMP1 to the surface of Plasmodium falciparum-infected erythrocytes. Mol Microbiol. 2008;68(5):1300–14.

44. Cooke BM, Buckingham DW, Glenister FK, Fernandez KM, Bannister LH, Marti M, et al. A Maurer’s cleft-associated protein is essential for expression of the major malaria virulence antigen on the surface of infected red blood cells. J Cell Biol. 2006;172(6):899–908.

45. Pachlatko E, Rusch S, Muller A, Hemphill A, Tilley L, Hanssen E, et al. MAHRP2, an exported protein of Plasmodium falciparum, is an essential component of Maurer’s cleft tethers. Mol Microbiol. 2010;77(5):1136–52.

46. Hanssen E, Sougrat R, Frankland S, Deed S, Klonis N, Lippincott-Schwartz J, et al. Electron tomography of the Maurer’s cleft organelles of Plasmodium falciparum-infected erythrocytes reveals novel structural features. Mol Microbiol. 2008;67(4):703–18.

47. Hirtzlin J, Farber PM, Franklin RM, Bell A. Molecular and biochemical characterization of a Plasmodium falciparum cyclophilin containing a cleavable signal sequence. Eur J Biochem. 1995;232(3):765–72.

48. Mok S, Ashley EA, Ferreira PE, Zhu L, Lin Z, Yeo T, et al. Drug resistance. Population transcriptomics of human malaria parasites reveals the mechanism of artemisinin resistance. Science. 2015;347(6220):431–5.

49. Bell A, Roberts HC, Chappell LH. The antiparasite effects of cyclosporin A: possible drug targets and clinical applications. Gen Pharmacol. 1996;27(6):963–71.

50. Mideo N, Reece SE, Smith AL, Metcalf CJ. The Cinderella syndrome: why do malaria-infected cells burst at midnight? Trends Parasitol. 2013;29(1):10–6.

51. Adjalley SH, Scanfeld D, Kozlowski E, Llinas M, Fidock DA. Genome-wide transcriptome profiling reveals functional networks involving the Plasmodium falciparum drug resistance transporters PfCRT and PfMDR1. BMC Genomics. 2015;16:1090.

52. Smith LM, Motta FC, Chopra G, Moch JK, Nerem RR, Cummins B, et al. An intrinsic oscillator drives the blood stage cycle of the malaria parasite Plasmodium falciparum. Science. 2020;368(6492):754–9.

53. Rijo-Ferreira F, Acosta-Rodriguez VA, Abel JH, Kornblum I, Bento I, Kilaru G, et al. The malaria parasite has an intrinsic clock. Science. 2020;368(6492):746–53.

54. Venkatesan M, Amaratunga C, Campino S, Auburn S, Koch O, Lim P, et al. Using CF11 cellulose columns to inexpensively and effectively remove human DNA from Plasmodium falciparum-infected whole blood samples. Malaria journal. 2012;11:41.

55. Cranmer SL, Magowan C, Liang J, Coppel RL, Cooke BM. An alternative to serum for cultivation of Plasmodium falciparum in vitro. Trans R Soc Trop Med Hyg. 1997;91(3):363–5.

56. Bolger AM, Lohse M, Usadel B. Trimmomatic: a flexible trimmer for Illumina sequence data. Bioinformatics. 2014;30(15):2114–20.

57. Dobin A, Davis CA, Schlesinger F, Drenkow J, Zaleski C, Jha S, et al. STAR: ultrafast universal RNA-seq aligner. Bioinformatics. 2013;29(1):15–21.

58. Patro R, Duggal G, Love MI, Irizarry RA, Kingsford C. Salmon provides fast and bias-aware quantification of transcript expression. Nat Methods. 2017;14(4):417–9.

59. Soneson C, Love MI, Robinson MD. Differential analyses for RNA-seq: transcript-level estimates improve gene-level inferences. F1000Res. 2015;4:1521.

60. Love MI, Huber W, Anders S. Moderated estimation of fold change and dispersion for RNA-seq data with DESeq2. Genome Biol. 2014;15(12):550.

61. Tavenard R FJ, Vandewiele G, Divo F, Androz G, Holtz C, Payne M, Yurchak R, Russwurm M, Kolar K, Woods E. tslearn: A machine learning toolkit dedicated to time-series data 2017.

62. Lopez-Barragan MJ, Lemieux J, Quinones M, Williamson KC, Molina-Cruz A, Cui K, et al. Directional gene expression and antisense transcripts in sexual and asexual stages of Plasmodium falciparum. BMC Genomics. 2011;12:587.

