## Supplemental Text, Figures, Tables for "Impact of sickle cell trait hemoglobin on the intraerythrocytic transcriptional program of *Plasmodium falciparum*"

### Supplemental Figures

**Fig S1**. Reads mapped per library of *in vitro* time series (A) or of in vivo samples (B).

**Fig S2**. Peak gene expression of *in vitro* time series.

**Fig S3**. Peak shifts of mono-peaking genes in 3D7 (A) or FUP (B) parasites.

**Fig S4**. Dynamic time warping scores by transcript and GO term.

**Fig. S5**. Stage-specific gene expression in *in vitro* time series.

**Fig. S6**. Expression of stage-specific genes *in vivo*.

**Fig. S7**. qPCR and RNA-seq measurements of *cyp19b*.

### Supplemental Tables

**Supplementary Table 1:** Genes differentially expressed in HbAS in both 3D7 and FUP.

**Supplementary Table 2:** Genes differentially expressed in HbAS *in vivo* samples.

### Supplemental Text

#### Time series experiments

Parasites of either 3D7 or FUP strain were expanded in culture with intermittent sorbitol synchronization to attain 12mL of infected red blood cells (iRBCs) at 2% parasitemia. To prepare early parasites that were between 0 and 3 hpi rings for our time series experiments, we adapted procedures from the Ring-stage Survival Assay (RSA) to isolate schizont stages using Percoll (1). Briefly, once the proportion of mature schizonts (each harboring between 10 and 12 nuclei) exceeded 0.5% of all RBCs in culture, the iRBCs were pelleted and resuspended in heparinized RPMI pre-warmed to 37°C. After a 15-minute incubation at 37°C, the culture was then gently layered onto a 75% Percoll solution, and centrifuged at 1,000g for 15 minutes. The intermediate phase containing mature schizonts was transferred to new tubes and washed with RPMI pre-warmed to 37°C. The schizonts were pooled in 2mL of ACM, and evenly distributed among 4 different flasks containing 1.8mL of red cells from four donors (two HbAA and two HbAS). These cultures were incubated for 3 hours to allow infection of donor RBCs and then treated with sorbitol to eliminate remaining schizonts and enrich for newly-infected donor RBCs. The resulting 1.8mL pellet of 0-3 hpi iRBCs from each donor was resuspended in 15mL of ACM, and 500µL was aliquoted into three 12-well plates containing 3.5mL of pre-warmed ACM per well, with each well containing 50µL of iRBCs. At each time point, two wells were harvested per donor. The first time point was taken immediately after the samples had been distributed, and each subsequent sample was taken at three-hour intervals. Harvested iRBCs were pelleted and resuspended in 1mL of TRIzol (Thermo Fisher). 30µL of suspended iRBCs was set aside at each time point for light microscopy using standard Giemsa staining. These slide readings to assess parasite maturation over the time course were performed by readers masked to red cell type and hpi of collection.

**Library preparation**

Samples from *in vitro* time series were homogenized in 1mL of TRIzol by passing them through a 0.6mm blunt needle coupled to a 1mL syringe 5 times, pelleted, and the supernatant was mixed with 200μL of chloroform. After centrifugation, the aqueous phase containing total cellular RNA was mixed with an equal volume of RNase-free 70% ethanol, and subsequently purified with the RNeasy Mini Kit (Qiagen). Samples were treated with DNase on-column, and RNA was eluted from the RNeasy mini column with 30μL of RNase-free water, quantified by Qubit High Sensitivity RNA Assay (Thermo Fisher Scientific), and stored at -80°C. For samples from the field study, we first added 0.75mL of TRIzol LS Reagent (Thermo Fisher) per 0.25mL of sample, and subsequently extracted RNA as described above.

Total RNA was submitted to the Duke Sequencing and Genomic Technologies Shared Resource, and libraries were prepared with the Kapa Stranded mRNA-seq library prep kit. Libraries from each of the two time series experiments (n=64/experiment) were sequenced on a full flow cell on the NovaSeq 6000 S2 platform with 150 base pair (bp) paired-end reads. Field study samples (n=32) were sequenced on a full flow cell of the NovaSeq 6000 S1 platform with 50bp paired-end reads.

**Transcriptome quantification**

Reads were assessed for quality with FastQC, and trimmed of adapter sequences and quality-filtered with Trimmomatic (2). Human reads were first depleted by aligning samples to the human genome (GRCh38.p13) with STAR (3), and the unmapped reads - which contained parasite transcripts - were saved in new fastq files. These parasite reads were quantified with Salmon (4) using the *Plasmodium falciparum* 3D7 transcriptome, assembly version ASM276v2. Quantification files of parasite gene expression were imported and summarized with the *tximport* package in R (5). Unnormalized read counts were prepared as input for DESeq2 and normalized abundance values expressed as transcripts per million (TPM) were prepared for other downstream analyses. Principal component analysis data was generated in R with the plotPCA function from DESeq2 after applying the variance-stabilizing transformation to count data. Data were then plotted with ggplot2. For heatmap visualization, TPMs were normalized around the mean for each transcript over the time series and depicted as a *z* score of standard deviations from the mean.

Scripts for RNA-seq read preparation and quantification are available on our GitHub page.

**Peak shift analysis**

Analyses of the shift of transcript peak were conducted by first identifying for each transcript in each time series the timepoint at which the transcript achieved its maximum expression, and then computing for each transcript the difference between each time series of the transcript's peak. All transcripts were included that exceeded 5 TPM at any time point over the time series for 3D7 (n= 4842 transcripts) or FUP (n=4657). For comparisons between ß-globin genotypes, HbAA samples served as the subtrahend and HbAS samples as the minuend. The difference was then expressed as a transcript's peak shift between samples, and, therefore, negative values represented a delay in a transcript's peak in HbAS parasites, and positive values an earlier expression peak in HbAS parasites. Peak shifts were also performed between HbAA controls, and served as control distributions against which to compare the HbAS shifts. To avoid skewing the results with large peak shifts arising from transcripts that have similar amplitudes of expression at the earliest and latest time points in the IDC, we restricted the shift window to +/- 42 hours. After removing these transcripts, the total number of genes under consideration for peak shift analysis was 4636 in 3D7, and 4559 in FUP.

We repeated this approach using only transcripts with a single peak over the time series. To identify mono-peaking transcripts, we utilized the find_peaks function from the signal processing submodule in SciPy, allowing a minimum peak width of 1 to capture transcripts that peak at the first and final time points. Mono-peaking transcripts were identified for each HbAA sample per strain, and the intersection of these transcripts was used as the list of mono-peak transcripts for 3D7 (n=891) or FUP (n=1067). We again restricted the shift window to +/- 42 hours. After removing these transcripts, the total number of transcripts under consideration for peak shift analysis was 830 in 3D7, and 1044 in FUP. We compared the distributions of peak shift values between samples using Kruskal-Wallis one-way analysis of variance. See Jupyter Notebook "expressionPeakChanges.ipynb" on our GitHub for full workflow.

**Differential expression analysis**

All differential expression analyses were performed with the DESeq2 (6) R package using unnormalized read counts per transcript. Time series samples were grouped by patient ß-globin genotype, parasite strain, and time point for differential gene expression analysis. Transcripts were excluded from DESeq2 analysis if they did not exceed 10 reads cumulatively over all samples, leaving 5,484 for analysis. Transcripts were considered differentially expressed if, as determined by DESeq2 with a false-discovery rate set to 0.05, both their adjusted p-value $< 0.05$ and $|\log_2(HbAS/HbAA)| > \log_2(1.5)$. *In vivo* patient samples were grouped by genotype and parasite stage. Similar to our time series experiments, transcripts were considered differentially expressed if, as determined by DESeq2 with a false-discovery rate set to 0.1, both the adjusted p-value $< 0.05$ and $|\log_2(HbAS/HbAA)| > \log_2(1.5)$.

**Gene ontology enrichment analysis**

Gene ontology enrichment analysis was performed with GOATOOLS (7). Gene IDs and associated GO IDs were downloaded from NCBI (ftp://ftp.ncbi.nlm.nih.gov/gene/DATA/gene2go.gz), and *P. falciparum's* annotations were extracted with the taxonomic identifier 36329. Using the transcripts identified as differentially expressed by DESeq2 according to the conditions described above as input, gene ontology terms were accepted for Benjamini-Hochberg adjusted p-values < 0.05. Differentially expressed transcripts shared between both 3D7 and FUP were organized by time point and separated into upregulated and downregulated groups. If the shared set of transcripts between strains was enriched for transcripts differentially expressed in the same direction (i.e., upregulated or downregulated in both strains) for the same GO term within one time point between strains, we considered that a shared response to HbAS. Some GO terms had significant overlap in their constituent transcripts. In these instances, we report the GO term that was the most specific in its functional description of the differentially expressed transcripts.

**Dynamic time warping analysis**

Dynamic time warping (DTW) of time series transcript expression data was computed with the tslearn toolkit (8). For each parasite strain, transcripts were excluded if transcript counts did not exceed 20 TPM in at least one time point for the entire time series analysis, while transcripts that did not exceed 5 TPM were excluded for ring-stage only time points. Time series were compared in a pairwise manner between replicates and ß-globin types. Owing to variation of expressed transcripts between strains, DTW scores were calculated for 3D7 and FUP separately. For each transcript, expression data were normalized to a mean of 0 with a standard deviation of 1 across the time series pair under investigation. This method generates scores for each transcript based on the overall similarity of their expression. The optimal alignment between two transcript expression series is displayed in the “warping path” through a matrix of Euclidean distances. Using a dynamic programming method, the path that produces the lowest cumulative score from this matrix is selected. Therefore, transcripts with similar expression profiles between two time series have low DTW scores, and the warping path lies near the diagonal of the matrix, while those that are expressed at varying time points between two time series have high DTW scores, and the warping path will likely deviate from the diagonal of the matrix. We did not constrain the search space or allowable slope of the warping path, so every point off of the diagonal could be considered to derive the optimal path between two time series for each transcript. For more information on this implementation, see the tslearn documentation (https://tslearn.readthedocs.io/en/stable/). When comparing individual transcript scores between HbAA and HbAS iRBCs, we considered as temporally-dysregulated transcripts with mean DTW scores across ß-globin replicates that exceeded two standard deviations above the mean DTW scores of all transcripts in HbAA versus HbAS comparisons. All analyses and data are available on our GitHub page (link). See Jupyter Notebook “DTW.ipynb” on our GitHub for full workflow.

**TAKT analysis**

The Temporal Alignment Kendall-Tau algorithm (TAKT) (9) was used to identify transcripts within each parasite that had similar dynamics but were temporally shifted during the ring stage between β-globin types. The TAKT algorithm allows for an assessment on the degree of dynamic similarity between two transcript abundance profiles by generating measures of curve-shape similarity and significance scores. The similarity measures are produced via the Kendal Tau (KT) dissimilarity metric which is a measure of discordance in the “up” and “down” patterns of transcript abundances between two transcript abundance profiles. By shifting one of the transcript abundance profiles and holding the other fixed and repeating the KT metric, measures of dynamic similarity across different time shifts are obtained. The time shift associated with the lowest KT measure is then defined as the estimate of how far apart in time a transcript is expressed between two timeseries. The significance of the TAKT measure is determined by permuting the time points of the shifted transcript abundance profile to generate random transcript abundance curves and computing the TAKT measure of dissimilarity between each random curve and the fixed transcript abundance profile. The empirical p-value is then the fraction of pairs whose dissimilarity is not larger than the dissimilarity between the true transcript abundance profiles. A more detailed description of the TAKT algorithm can be found in (9). We first truncated all timeseries datasets to only contain transcript abundances measured during the ring stage of the respective parasite (3D7: 0-24 hours; FUP: 0-18 hours). Measures of similarity and associated significance scores were produced for a gene between similar β-globin type replicates and across different β-globin type replicates within a parasite using TAKT. The similarity scores produced by TAKT are independent of transcript abundance values, which allowed us to only measure the similarity of transcript dynamics, rather than the similarity between expression levels. By focusing on only the ring stage portion of the data, we were able to calculate the exact sampling distribution through exhaustive enumeration of the ring-stage timepoints, thus allowing us to calculate the exact p-values of the similarity measures.

We next queried the TAKT results to find transcripts that had similar dynamics across all β-globin types within each parasite but displayed minimal shifts between HbAA replicates and large shifts between HbAA and HbAS replicates. A p-value of ≤ 0.05 was used to identify transcripts with similar dynamics across all β-globin types within each parasite. We examined three different ranges of time shifts for each transcript, +/- 3, 6, or 9 hours. For example, a gene was determined similarly phased in HbAA replicates but not in HbAS replicates if the transcript in one HbAA replicate was within +/- 3 hours of its other HbAA replicate and both its HbAS replicates were phased > +/-3 hours from each HbAA replicate. All analyses and data are available on our GitHub page (link). See Jupyter Notebooks “TAKT_analysis.ipynb” and “DTW.ipynb” on our GitHub for workflow.

**Stage classification of *in vivo* parasites**

Stage classification of *in vivo* parasites was determined by first inspecting normalized expression heatmaps sorted by peak expression. Based on morphology assessed by microscopy, we identified the set of transcripts in each strain from our *in vitro* time series that peaked within each stage of the IDC. From this, transcripts for rings, trophozoites, and schizont stage parasites were determined for each strain. We took the intersection of the 3D7 and FUP sets to identify the stage-specific transcripts that are shared between both strains of *P. falciparum.* Finally, we selected transcripts that were highly expressed at their peak, discarding those that did not exceed 1,000 TPM. See Jupyter Notebook “Genes_by_stage.ipynb” for full workflow. We also estimated parasite stage with a previously published mixture model (10). Our count data was converted to RPKM and log2-transformed to match the staged expression data (11) used for stage estimation.

**All transcripts - peak shift analysis**

The proportion of all transcripts that peaked at the same timepoint in HbAA and HbAS RBCs was 21.4% in 3D7 (1036/4636) and 22.9% (1066/4559) in FUP parasites, while the proportion of transcripts that peaked in HbAS RBCs within 3 hours or less of their peak expression time in HbAA RBCs was 60% in 3D7 (2926/4636) and 56.4% in FUP (2626/4559) parasites.  We observed minimal background variance in transcript peaks when comparing parasites growing HbAA RBCs: with peak shifts of 0.56 hours (3D7) and 0.78 hours (FUP). When compared to these parasite growing in HbAA, mean peak shifts indicated slight delay in in HbAS RBCs for both 3D7 (-1.91 hours) and FUP (-1.16 hours) parasites), though these differences were not significantly different by Kruskal-Wallis one-way ANOVA (3D7: p = 1.0; FUP: p = 1.0.

### Transcriptome quantification

Of the 128 RNA-seq libraries generated in the time-series experiments, an average of 24,285,500 fragments per library mapped to the *P. falciparum* 3D7 transcriptome (assembly version ASM276v2). The mean (range) number of fragments mapped per sample was 27,195,400 (10,695,730 -- 46,016,550) for 3D7 parasites and 21,375,600 (6,723,168 -- 39,655,690) for FUP parasites. As parasites matured during the IDC, the number of mapped fragments increased (**Figure S1**).

In the 32 libraries generated from freshly-collected parasites in the field study, all libraries exceeded one million fragments mapped, achieving sufficient coverage across the *P. falciparum* transcriptome in all 32 RNA-seq libraries (**Figure S1B**). The mean (range) number of fragments mapped to the 3D7 transcriptome was 10,390,430 (1,017,756-36,409,570) for parasites in children with HbAA and 7,749,058 (1,696,908-27,364,260) for those with HbAS.

### Transcriptional synchrony by peak-shift of transcripts

The proportion of all transcripts that peaked at the same timepoint in HbAA and HbAS iRBCs across all pairwise comparisons was 24.8% in 3D7 (206/830) and 25.5% in FUP (266/1044); similarly, the proportion of transcripts that peaked in HbAS iRBCs within 3 hours or less of their peak expression in HbAA iRBCs was 73.4% in 3D7 and 66% in FUP parasites.

Similar to the results of peak-shift analysis of single-peaking transcripts, we repeated these analyses among all transcripts without restricting to those with a single transcriptional peak, and also observed minimal shifts of peak transcript expression in HbAS RBCs.

**Intersection of aberrant transcripts in HbAS across analytic approaches**

In our *in vitro* experiments, we observed differential expression during at least 1 timepoint for 2499 3D7 (45.6%) and 3215 FUP (58.6%) transcripts; 1683 of these were shared between both strains. DTW analyses identified fewer aberrantly expressed transcripts compared to differential expression analysis, with only 155 3D7 (3.2%) and 162 FUP (3.48%) parasite transcripts temporally dysregulated by DTW; 37 (0.8%) of these were shared between strains.

Over half of the transcripts (3D7: 70.3%; FUP: 58.6%) that were identified as temporally dysregulated by DTW were also identified as differentially expressed within strains.

A

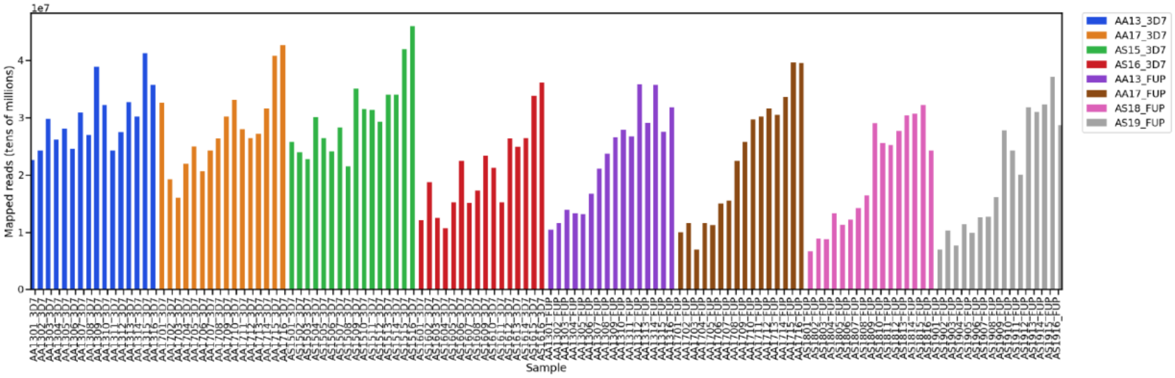


B

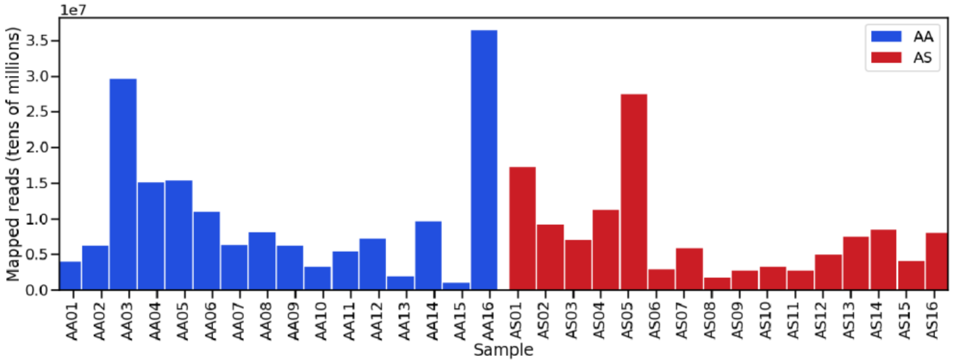


**Fig S1. Reads mapped per library of *in vitro* time series (A) or of in vivo samples (B). A:** Number of fragments mapped for each library from 3D7 and FUP time course experiments. Bars are colored by time series sample. B: Number of fragments mapped for each parasite transcriptome from Malian children. Bars are colored by β-globin genotype.

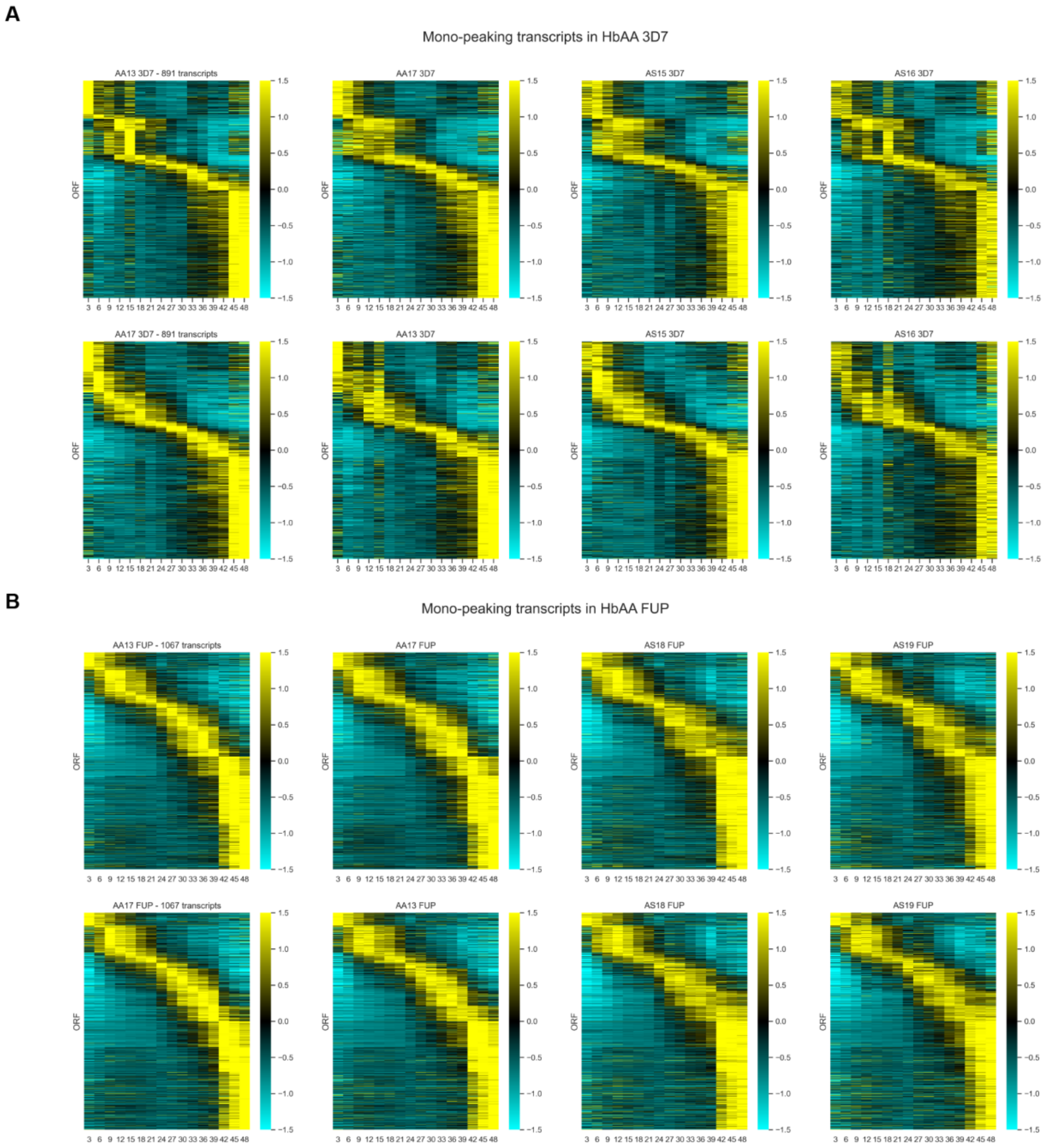


**Fig S2. Peak gene expression of *in vitro* time series.** Ad **(A)** Peak expression time of the subsets of mono-peaking parasite transcripts in HbAA and HbAS iRBCs for 3D7 parasites. For each row, transcripts are ordered vertically by the peak expression time in the first or second HbAA replicate. **(B)** Peak expression time of the subsets of mono-peaking parasite transcripts in HbAA and HbAS iRBCs for FUP parasites. For each row, transcripts are ordered vertically by the peak expression time in the first or second HbAA replicate. The columns along the *x*-axis depict the normalized expression values of each transcript per time point. Expression values are normalized around the mean for each transcript over the time series and depicted as a *z* score of standard deviations from the mean. Values range between -1.5 (cyan) and 1.5 (yellow).

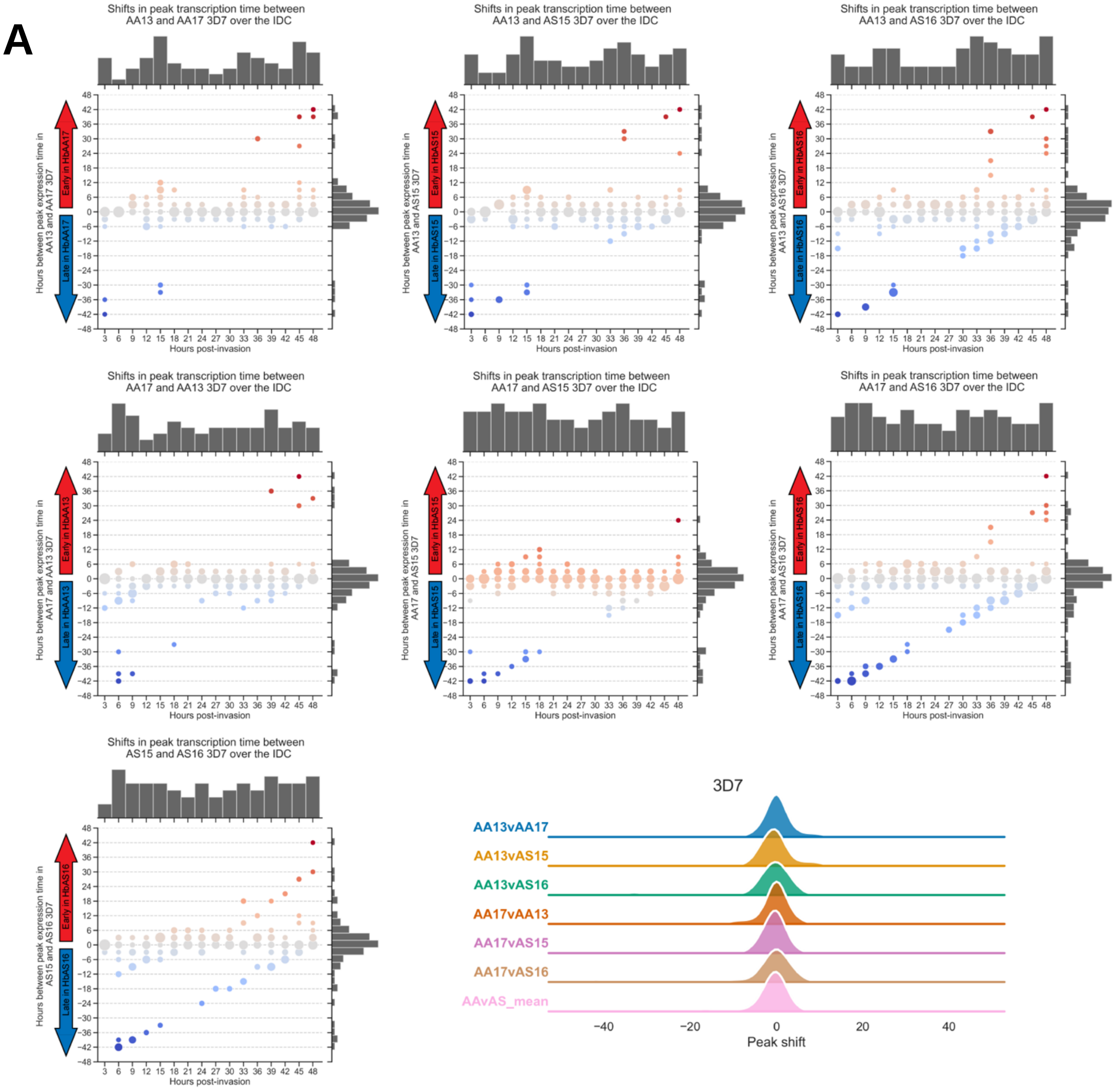
A
Shifts in peak transcription time between AA13 and AA17 3D7 over the IDC
Shifts in peak transcription time between AA13 and AS15 3D7 over the IDC
Shifts in peak transcription time between AA13 and AS16 3D7 over the IDC
Shifts in peak transcription time between AA17 and AA13 3D7 over the IDC
Shifts in peak transcription time between AA17 and AS15 3D7 over the IDC
Shifts in peak transcription time between AA17 and AS16 3D7 over the IDC
Shifts in peak transcription time between AS15 and AS16 3D7 over the IDC
Hours between peak expression time in AA13 and AA17 3D7
Hours between peak expression time in AA13 and AS15 3D7
Hours between peak expression time in AA13 and AS16 3D7
Hours between peak expression time in AA17 and AA13 3D7
Hours between peak expression time in AA17 and AS15 3D7
Hours between peak expression time in AA17 and AS16 3D7
Hours between peak expression time in AS15 and AS16 3D7
Early in HbAA17
Late in HbAA17
Early in HbAS15
Late in HbAS15
Early in HbAS16
Late in HbAS16
Early in HbAA13
Late in HbAA13
Hours post-invasion
3D7
AA13vAA17
AA13vAS15
AA13vAS16
AA17vAA13
AA17vAS15
AA17vAS16
AAvAS_mean
Peak shift

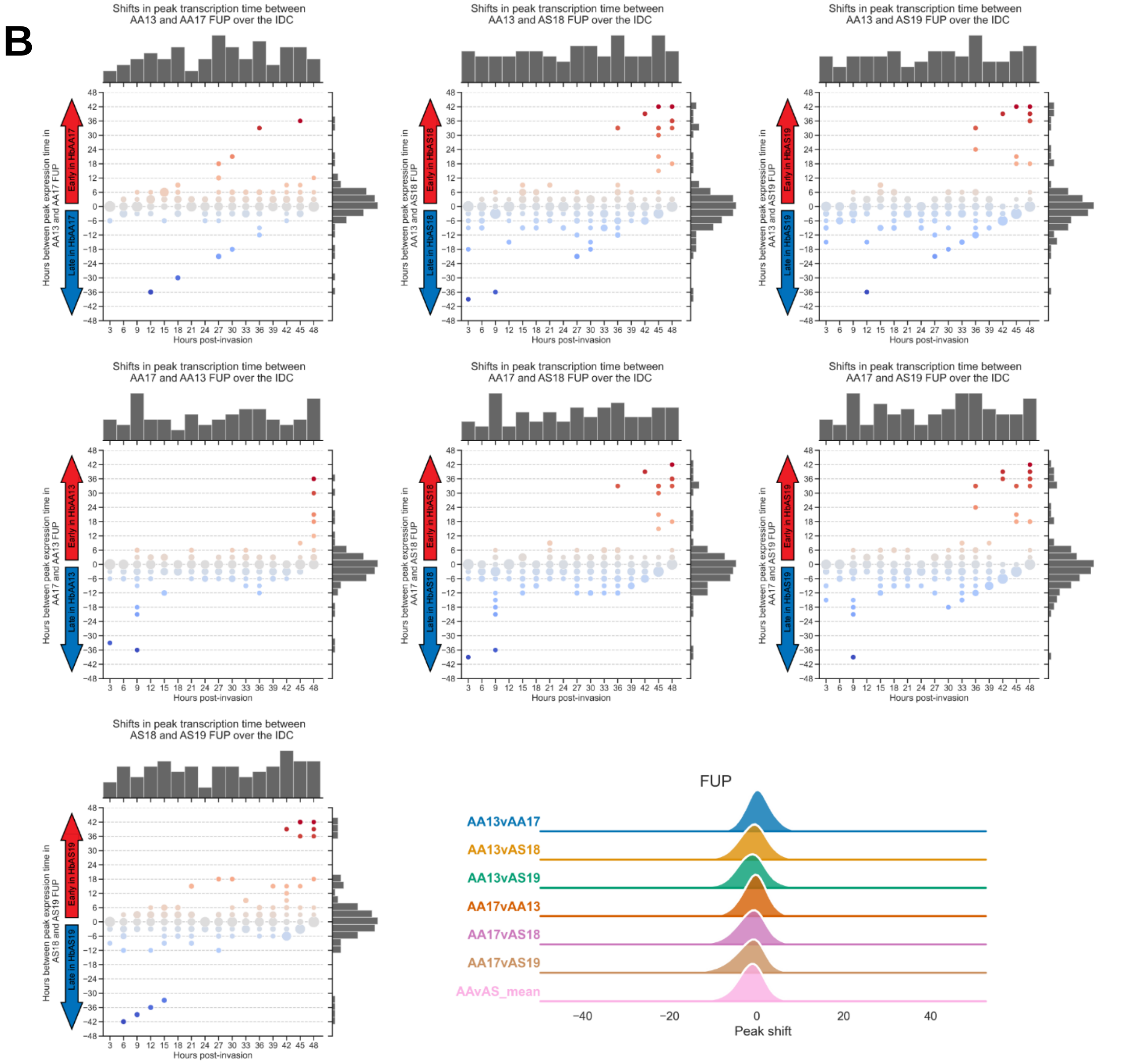


**Fig S3. Peak shifts of mono-peaking genes in 3D7 (A) or FUP (B) parasites.**

Temporal shifts for peak expression timepoint for the subset of mono-peaking parasite transcripts between each sample. For each transcript, the time between its expression peak was calculated between samples. Each *x*-axis category indicates the number of transcripts that normally peak at this timepoint in the first listed sample, and the y-values indicate the distributions of the shifts of the peaks of these transcripts in the second sample. Circles are sized relative to the number of transcripts with that y-value. Top histogram indicates the number of parasite transcripts that peak at each *x*-value timepoint, and right histogram indicates the number of transcripts with peak shifts of each *y*-value. The distribution of peak shifts (displayed along the right *y*-axis) is centered around zero, indicating that most transcripts peak at the same time in the time series samples.

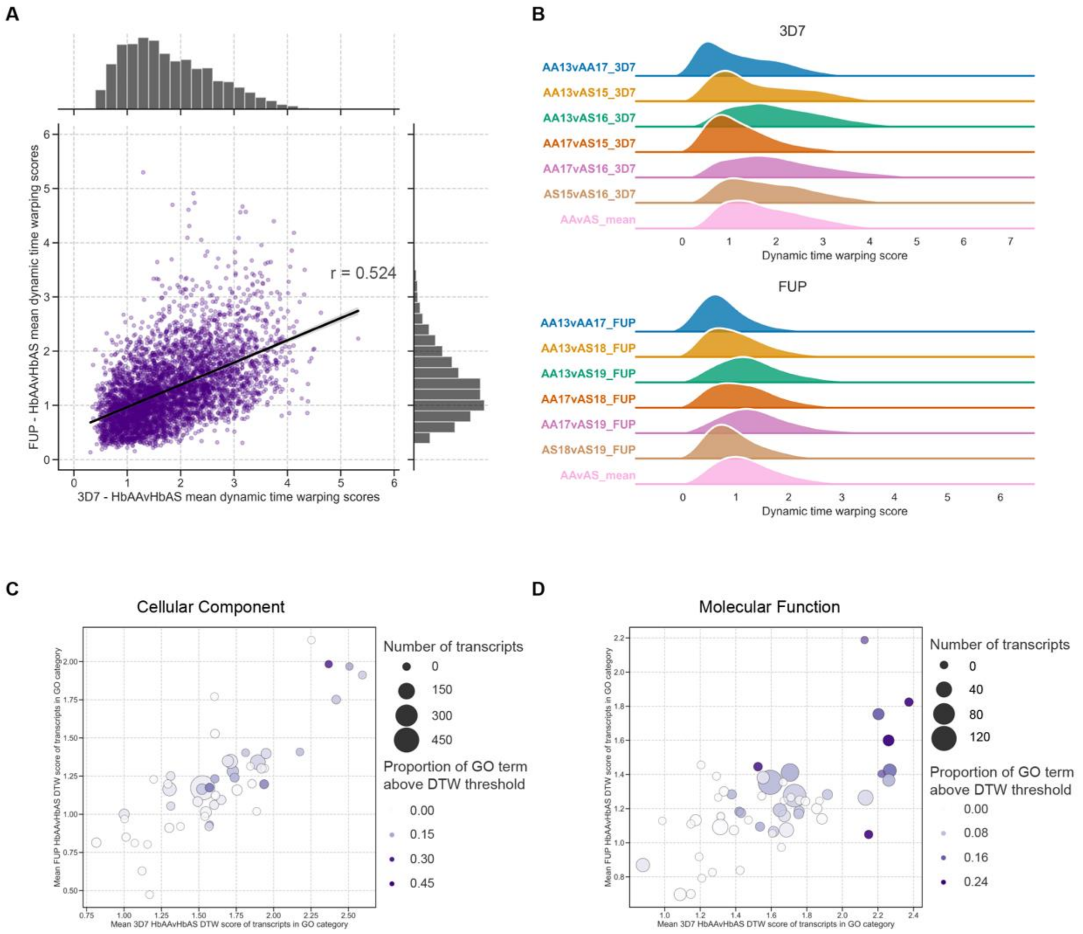


**Fig S4. Dynamic time warping scores by transcript and GO term. (A)** Scatterplot of dynamic time warping (DTW) scores in time series data of individual transcripts in HbAA compared to HbAS samples in 3D7 (*x*-axis) and FUP (*y*-axis) parasites. For each strain, scores were computed pairwise between HbAA and HbAS samples and averaged. Transcripts with dynamic time warping scores greater than two standard deviations from the overall mean (points to the right of the red line for 3D7, and points above the blue line for FUP) were considered temporally dysregulated. Histograms indicate overall distribution of transcript DTW scores in either 3D7 (top) or FUP (right). Line indicates Pearson correlation coefficient. **(B)** Distributions of DTW scores for each pairwise comparison and averaged DTW scores across β-globin replicates. The distributions of DTW scores across all comparisons were not statistically different (Kruskal-Wallis one-way ANOVA – 3D7: p=1.0; FUP: p=1.0). **(C)** Scatterplot of mean DTW scores within Cellular Component GO terms that are enriched for temporally-dysregulated transcripts in HbAS iRBCs in both 3D7 (*x*-axis) and FUP (*y*-axis) parasites. **(D)** Scatterplot of mean DTW scores within Molecular Function GO terms that are enriched for temporally-dysregulated transcripts in HbAS iRBCs in both 3D7 (*x*-axis) and FUP (*y*-axis) parasites. In **(C)** and **(D)**, color indicates GO term, shading indicates hpi (darker = later hpi), and size of point indicates the proportion of transcripts within the GO term that are temporally-dysregulated as defined by a DTW score greater than 2 standard deviations from the overall mean of DTW scores (points beyond the dashed lines in **Figure 5A**). Between 3D7 and FUP, GO-level DTW scores were highly correlated for Biological Process (r = 0.779, p-value < 0.001), Cellular Component (r = 0.781, p-value < 0.001) and Molecular Function (r = 0.656, p-value < 0.001) categories, indicating that HbAS impacts the timing of transcription of the same functional groups across the two *P. falciparum* strains.

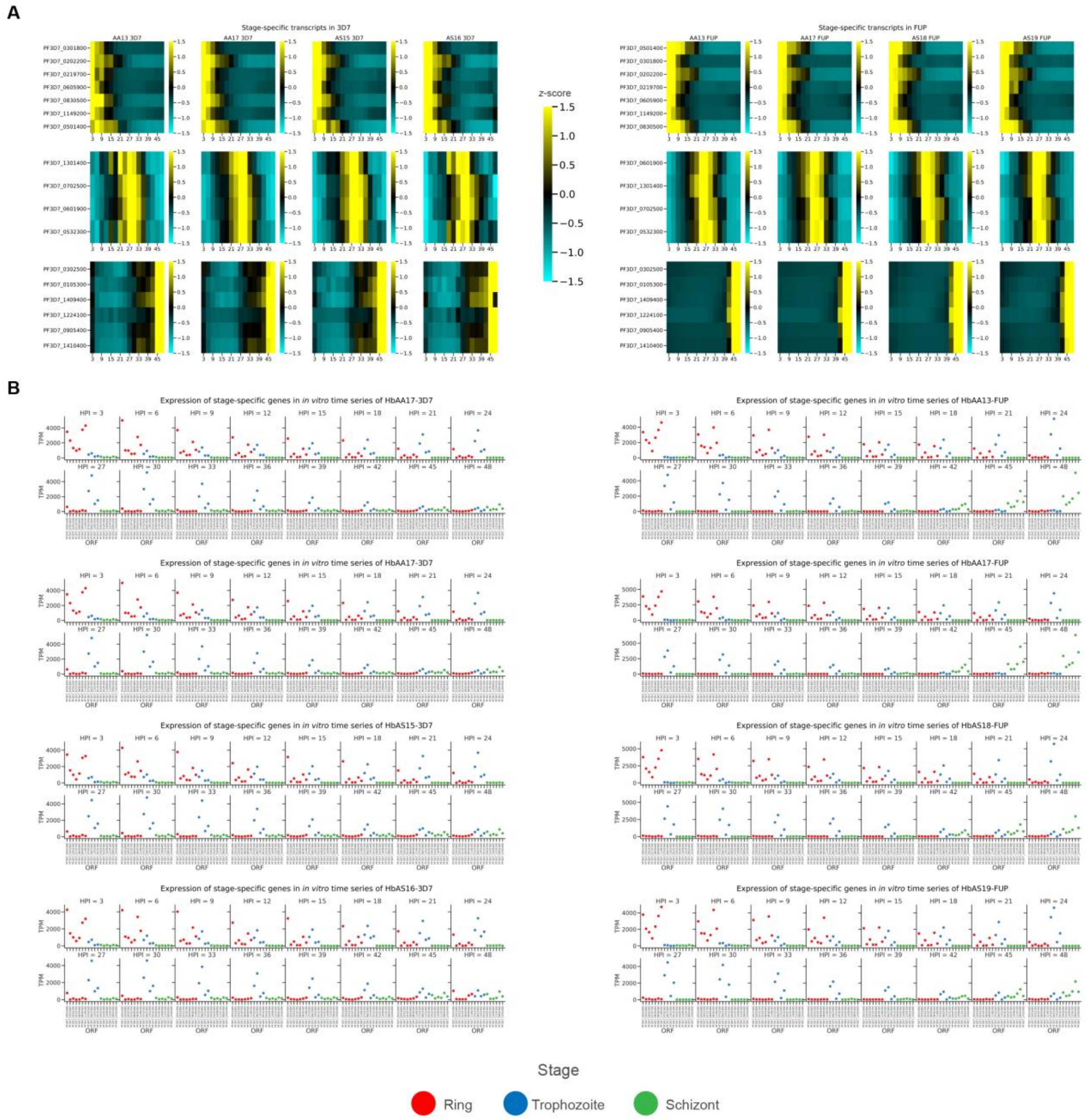


**Fig. S5. Stage-specific gene expression in *in vitro* time series. (A)** Heatmaps of transcripts that are expressed in a stage-specific manner across 3D7 and FUP time series experiments. Transcripts for each parasite strain are order vertically by their peak expression time in HbAA iRBCs. The ordering of peak expression in HbAS samples is not conserved in temporally dysregulated transcripts identified by DTW. Transcripts are ordered by peak expression time in HbAA samples and displayed as a *z* score of standard deviations from the mean. **(B)** Normalized expression of stage-specific transcripts for each time series sample. Points are colored by stage assignments.

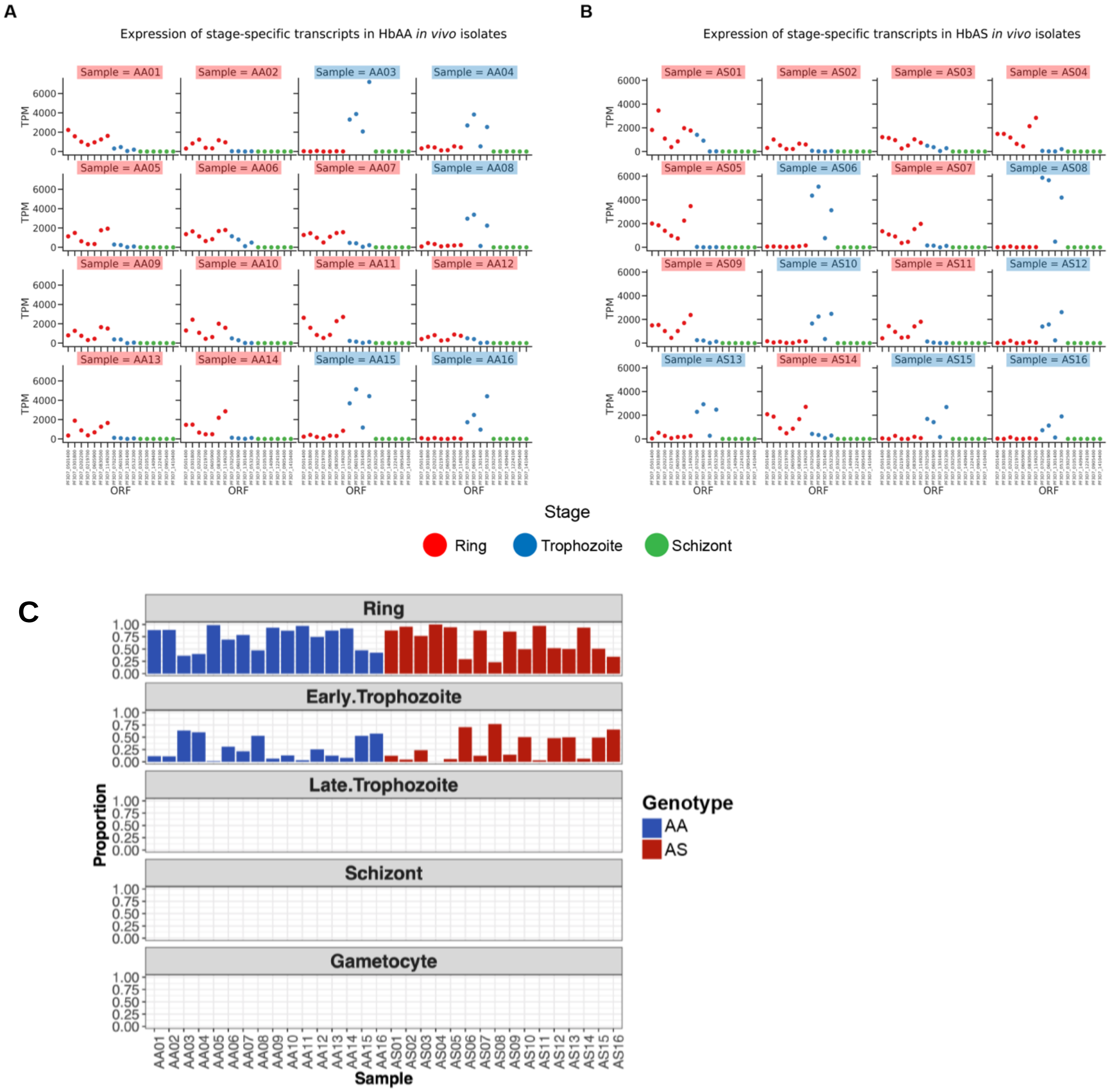


**Fig. S6. Expression of stage-specific genes *in vivo*. (A)** Normalized expression of stage-specific transcripts in each sample from HbAA and **(B)** HbAS Malian children. Points are colored by stage assignment, and sample names are colored by the stage classification assigned according to abundance of stage-specific transcripts. **(C)** Estimates of stage of each infection using an independent published mixture model from (10).

A

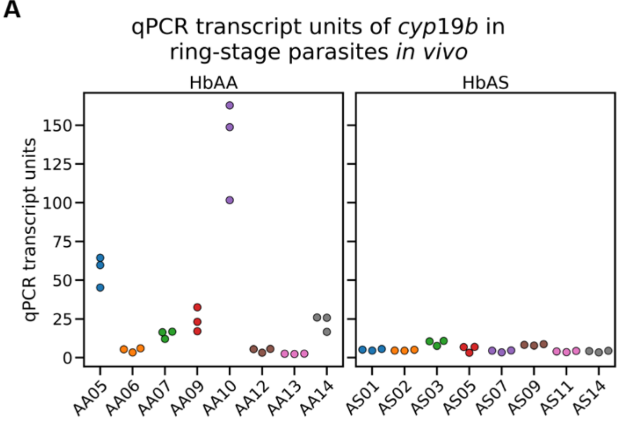


B

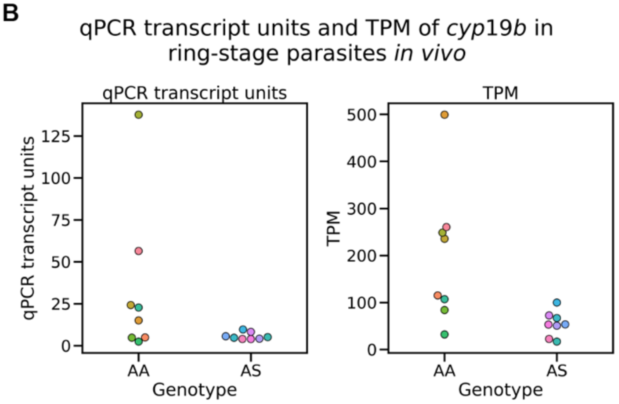


**Fig. S7. qPCR and RNA-seq measurements of *cyp19b*. (A)** Transcript units measured by qPCR of *cyp19b* in parasites from HbAA and HbAS patients. Each biological replicate was measured in triplicate. **(B)** Average qPCR transcript units from technical triplicates of each *in vivo* sample (left), and TPM values for each sample from RNA-seq (right).


1. Witowski B MD, Amaratunga C, Fairhurst RM. Ring-stage Survival Assays (RSA) to evaluate the in-vitro and ex-vivo susceptibility of Plasmodium falciparum to artemisinins. National Institutes of Health Procedure RSAv1. Institute Pasteur du Cambodge2013.
2. Bolger AM, Lohse M, Usadel B. Trimmomatic: a flexible trimmer for Illumina sequence data. Bioinformatics. 2014;30(15):2114-20.
3. Dobin A, Davis CA, Schlesinger F, Drenkow J, Zaleski C, Jha S, et al. STAR: ultrafast universal RNA-seq aligner. Bioinformatics. 2013;29(1):15-21.
4. Patro R, Duggal G, Love MI, Irizarry RA, Kingsford C. Salmon provides fast and bias-aware quantification of transcript expression. Nat Methods. 2017;14(4):417-9.

5. Soneson C, Love MI, Robinson MD. Differential analyses for RNA-seq: transcript-level estimates improve gene-level inferences. F1000Res. 2015;4:1521.
6. Love MI, Huber W, Anders S. Moderated estimation of fold change and dispersion for RNA-seq data with DESeq2. Genome Biol. 2014;15(12):550.
7. Klopfenstein DV, Zhang L, Pedersen BS, Ramirez F, Warwick Vesztrocy A, Naldi A, et al. GOATOOLS: A Python library for Gene Ontology analyses. Sci Rep. 2018;8(1):10872.
8. Tavenard R FJ, Vandewiele G, Divo F, Androz G, Holtz C, Payne M, Yurchak R, Russwurm M, Kolar K, Woods E. tslearn: A machine learning toolkit dedicated to time-series data 2017.
9. Kelliher CM, Foster MW, Motta FC, Deckard A, Soderblom EJ, Moseley MA, et al. Layers of regulation of cell-cycle gene expression in the budding yeast Saccharomyces cerevisiae. Mol Biol Cell. 2018;29(22):2644-55.
10. Tonkin-Hill GQ, Trianty L, Noviyanti R, Nguyen HHT, Sebayang BF, Lampah DA, et al. The Plasmodium falciparum transcriptome in severe malaria reveals altered expression of genes involved in important processes including surface antigen-encoding var genes. PLoS Biol. 2018;16(3):e2004328.
11. Lopez-Barragan MJ, Lemieux J, Quinones M, Williamson KC, Molina-Cruz A, Cui K, et al. Directional gene expression and antisense transcripts in sexual and asexual stages of Plasmodium falciparum. BMC Genomics. 2011;12:587.
